# Coarse-grained molecular dynamics simulations of lipid-protein interactions in SLC4 proteins

**DOI:** 10.1101/2023.06.26.546592

**Authors:** Hristina R. Zhekova, Daniel P. Ramirez-Echemendía, Besian I. Sejdiu, Alexander Pushkin, D. Peter Tieleman, Ira Kurtz

## Abstract

The SLC4 family of secondary bicarbonate transporters is responsible for the transport of HCO_3_−, CO_3_^2−^, Cl^−^, Na^+^, K^+^, NH_3_ and H^+^ necessary for regulation of pH and ion homeostasis. They are widely expressed in numerous tissues throughout the body and function in different cell types with different membrane properties. Potential lipid roles in SLC4 function have been reported in experimental studies, focusing mostly on two members of the family: AE1 (Cl^−^/HCO_3_^−^ exchanger) and NBCe1 (Na^+^-CO_3_^2−^ cotransporter). Previous computational studies of the outward facing (OF) state of AE1 with model lipid membranes revealed enhanced protein-lipid interactions between cholesterol (CHOL) and phosphatidylinositol bisphosphate (PIP2). However, the protein-lipid interactions in other members of the family and other conformation states are still poorly understood and this precludes the detailed studies of a potential regulatory role for lipids in the SLC4 family. In this work, we performed multiple 50 µs coarse-grained molecular dynamics simulations on three members of the SLC4 family with different transport modes: AE1, NBCe1 and NDCBE (a Na^+^-CO_3_^2−^/Cl^−^ exchanger), in model HEK293 membranes consisting of CHOL, PIP2, phosphatidylcholine (POPC), phosphatidylethanolamine (POPE), phosphatidylserine (POPS), and sphingomyelin (POSM). The recently resolved inward-facing (IF) state of AE1 was also included in the simulations. Lipid-protein contact analysis of the simulated trajectories was performed with the ProLint server, which provides a multitude of visualization tools for illustration of areas of enhanced lipid-protein contact and identification of putative lipid binding sites within the protein matrix. We observed enrichment of CHOL and PIP2 around all proteins with subtle differences in their distribution depending on the protein type and conformation state. Putative binding sites were identified for CHOL, PIP2, POPC, and POSM in the three studied proteins and their potential roles in the SLC4 transport function, conformational transition and protein dimerization were discussed.

**Statement of significance:** The SLC4 protein family is involved in critical physiological processes like pH and blood pressure regulation and maintenance of ion homeostasis. Its members can be found in various tissues. A number of studies suggest possible lipid regulation of the SLC4 function. However, the protein-lipid interactions in the SLC4 family are still poorly understood. Here we make use of long coarse-grained molecular dynamics simulations to assess the protein-lipid interactions in three SLC4 proteins with different transport modes, AE1, NBCe1, and NDCBE. We identify putative lipid binding sites for several lipid types of potential mechanistic importance, discuss them in the framework of the known experimental data and provide a necessary basis for further studies on lipid regulation of SLC4 function.

## Introduction

Transporter proteins are involved in translocation of a large variety of substrates across cellular and organelle membranes. A plethora of studies has provided evidence for the critical role of membrane lipids and specific lipid-protein interactions on the transport properties of membrane proteins. The lipid effects on protein function are fairly diverse (1,2). Some examples include lipid-driven protein oligomerization and aggregation, state-specific modulation of transport, lipid mediated signaling, and specific sequestering of proteins into ordered/disordered membrane areas or areas characterized with positive/negative membrane curvature. Recent resolution of X-Ray and cryoEM structures with bound lipids coupled with molecular modelling allows assessment of specific lipid-protein interactions and their functional role at the molecular level. Nevertheless, our understanding of the role of the lipid membrane on protein function is still insufficient and critical experimental and computational data on lipid-protein interactions is lacking for multiple protein families, especially those featuring secondary transporters, which have been historically difficult to characterize with available structural methods.

The SLC4 family of bicarbonate and carbonate transporters is involved in multiple processes of physiological importance, such as pH and blood pressure regulation and maintenance of ion homeostasis (3-9). The family consists of 11 members that include Na^+^-anion symporters and both Na^+^-independent and Na^+^-dependent anion exchangers. Their presence in various cell types raises questions about the possible modulating role of lipid-protein interactions which may vary in the different cellular membrane environments. Several experimental studies have focused on the lipid dependence on SLC4 transport function. AE1 (SLC4A1, Anion exchanger 1, or Band 3 protein) is found predominantly in erythrocytes and α-intercalated cells of the renal collecting duct where it exchanges Cl^−^ for HCO_3_^−^ (7). Various studies on the effect of lipids on AE1 function have been done in erythrocytes, erythrocyte ghost membranes, and model monolayers and liposomes and have demonstrated dependence of AE1 aggregation and function on phosphatidylcholine (PC), phosphatidylserine (PS), sphingomyelin (SM), cholesterol (CHOL) and phosphatidylinositol (PI) lipids (10-17). This dependence has been explained either as a result of membrane rigidification in the presence of CHOL and saturated lipid tails from the SM, PS, and PC lipids or as allosteric inhibition of anion transport (16). Computational studies of AE1 dimers in model lipid bilayers showed aggregation of PIP2 lipids around the protein and preference for CHOL binding at the dimeric interface (18,19). NBCe1 (SLC4A4) is an electrogenic sodium-carbonate symporter present in the renal proximal tubule cells where it transports Na^+^-CO_3_^2−^ across the basolateral membrane (5,7,20). Patch-clamp measurements in HeLa cells and Xenopus oocytes of NBCe1-A, B, and C have provided evidence of activation of all three isoforms by phosphatidylinositol bisphosphate (PIP2), although NBCe1-A has higher apparent affinity for PIP2 than the other two isoforms which possess an auto-inhibitory domain in their N-terminus (N_t_) (21-23). Additionally, a putative binding site for PIP2 has been identified in the N_t_ domains of NBCe1-B and C, although the mechanism of PIP2 activation and transport modulation on the overall transport function of NBCe1 is still not understood (21).

Recent X-ray and cryoEM structures of the transmembrane domains of human AE1 (hAE1) (24), human NBCe1 (hNBCe1) (25), and rat NDCBE (rNDCBE, a Na^+^-CO_3_^2−^/Cl^−^ exchanger that combines features of AE1 and NBCe1) (26) in the outward facing (OF) state and structures of bovine AE1 in the OF and inward facing (IF) state (27) led to identification of putative anion and cation binding sites and ion and water permeation pathways within the protein matrix (26-28) and provided evidence of elevator-like transport mechanism in the SLC4 family and other proteins with similar architecture, i.e. the SLC23 and SLC26 families (29-31), the bacterial uracil transporter UraA (32), the fungal purine transporter UapA (33), and the plant borate transporter Bor1 (34). The new structures create opportunities for computational assessment of lipid-protein interactions in the SLC4 family, which can provide further details on the system-specific and state-specific lipid-protein interactions in this family. In the current work, we report results from multiple 50 µs long coarse-grained molecular dynamics simulations in a simplified model HEK293 membrane of the available homodimers of human AE1_OF-OF_, NBCe1, and rat NDCBE in outward facing state, and the newly reported bovine AE1_IF-IF_ homodimers which feature inward facing states. We observe both common and system-specific and state-specific lipid-protein interactions and identify putative binding sites for the lipids with most prominent protein contacts. Possible mechanistic implications of the lipid-protein interactions on the OF to IF conformational transition in the SLC4 family are discussed as well.

## Methods

### System setup

Coarse grained molecular dynamics simulations (CGMD) were performed on systems containing a single protein dimer in a simplified model bilayer mimicking the HEK293 plasma membrane, since the HEK293 cell line is one of the most common systems for functional measurements of SLC4 transport (25-28). The following systems were included in the study: human AE1 (AE1_OF-OF_), human NBCe1, and rat NDCBE, as homodimers in which the monomers were in the outward facing (OF) open state (OF-OF dimers) and bovine AE1 (AE1_IF-IF_) as homodimers with inward facing (IF) open monomers (IF-IF dimers). The corresponding PDB accession codes for the OF-OF dimers of AE1_OF-OF_, NBCe1, and NDCBE used for preparation of the simulated systems are 4YZF (24), 6CAA (25), and 7RTM (26), respectively. A recent NBCe1 structure of improved resolution (I.K. and A.P, unpublished data, 2022) was used for modelling of the large extracellular loop 3 and any missing residues in this loop were further modelled by Modeller 10.3 (35). This loop is mostly positioned in the extracellular solution and is not involved in lipid-protein interactions, as evident from our CGMD simulations (see below). The initial coordinates of the AE1_IF-IF_ dimers were taken from PDB 8D9N (27) and optimized further with the Molecular Dynamics Flexible Fitting (MDFF) (36,37) method as described in Ref.(27). The final AE1 _IF-IF_ model used in our simulations includes also the H1 helix, which is missing in the deposited AE1_IF-IF_ homodimer of 8D9N. The proteins were converted into a coarse-grained representation, using martinize.py v2.6 with the Martini v2.2 force field (38) and elnedyn22 (39) elastic constraints imposed on the backbone atoms. Two relevant disulfide bonds between Cys residues in EL3 in rNDCBE (C638-C672 and C636-C684) and hNBCe1 (C585-C630 and C583-C642) were introduced during this process. Afterwards, each dimer was embedded in a lipid membrane and solubilized in a 0.15 NaCl water solution with insane.py (40). The resultant periodic box had final dimensions of 20 × 20 × 30 nm. The model HEK293 membrane bilayer featured the following composition: upper leaflet: POPC:POPE:POSM:POPS:PIP2:CHOL 33:4:7:0:0:13; lower leaflet: POPC:POPE:POSM:POPS:PIP2:CHOL 14:14:3:20:5:12. The POP2 topology and parameters were used for the PIP2(3,4) representation in the CGMD simulations. The martinize.py and insane.py scripts were downloaded from the MARTINI website: http://cgmartini.nl/index.php/tools2/proteins-and-bilayers.

### Coarse grained MD simulations

Three replicas per protein dimer were prepared as described above and submitted to 50 µs long coarse-grained MD simulations with the Martini v2.2 (38) force field and Gromacs version 2020.6 (41). A short energy minimization step with the steepest descent algorithm, followed by a stepwise equilibration process with gradual release of constraints on the protein atoms were done before the production runs, as described previously (42). The production runs were done at 310 K temperature and 1 bar pressure maintained with the v-rescale thermostat (time coupling τ = 1.0 ps) and semi-isotropic Parrinello-Rahman barostat (compressibility = 3×10^−4^ bar, relaxation time τ = 12 ps), respectively. Electrostatic effects were evaluated via the reaction-field scheme with r_Coulomb_ = 1.1 nm, r_VdW_ = 1.1 nm, and ε_r_ = 15. A 20fs time step with LINCS constraints for the side-chain beads was adopted for the simulations.

### Trajectory analysis

The last 20 µs of the generated 50 µs trajectories were used for analysis of lipid-protein interactions with the ProLint server (43) developed in the Tieleman lab and in-house scripts for depletion and enrichment index calculations. Lipids reached equilibration around the protein within the first 10 µs of simulation. Inspection of trajectories and produced results (metric graphs, heat maps, density plots, membrane thickness, etc.) show consistent behavior for all studied lipids in all studied replicas and in the monomers of the homodimers. The lipid-protein contact analysis makes use of a 0.6 nm cutoff radius, unless stated otherwise. This cutoff radius covers the first coordination shell for important lipid types, such as PIP2 and CHOL, as demonstrated in previous computational studies of GPCR-lipid interactions (42). The depletion-enrichment index calculations were done as described previously (42).

## Results

### Reorganization of different lipid types around the protein matrix from lipid density distribution and lipid depletion and enrichment analysis

Figure 1 (top) shows density distribution plots for the six studied lipids around the proteins in selected representative replicas. A full set of figures from all replicas is included in Figure S1. The contrast between dark/light areas on the map serve as a semi-quantitative representation of the lipid density difference in different areas of the density map (i.e. darker areas signify lower lipid density than the bulk density and lighter areas signify higher lipid density than the bulk density for a specific lipid in the system). There is evident increase of density (brighter spots around the protein matrix) of CHOL and PIP2 around all proteins, regardless of the conformation state. Increased POSM density is found at the dimerization interface of hNBCe1 and AE1_IF-IF_. The remaining three lipids (POPC, POPE, and POPS) do not feature substantial density difference between bulk and protein-adjacent areas. To quantify the depletion or enrichment of the different lipid types around the protein matrix, with respect to the average lipid distribution in the bulk, we performed calculations of depletion-enrichment indices (DEI) for all studied systems, using a cutoff value of 0.6 nm. The DEI averaged over each 3 replicas are presented in Figure 1 (bottom). A number larger or smaller than 1 signifies enrichment or depletion, respectively, of lipids of specific type around the protein matrix. Consistent with the density distribution plots in Figure 1 (top) all studied systems demonstrate considerable enrichment of PIP2 lipids in the area adjacent to the protein. A smaller, but nevertheless significant enrichment of CHOL is also evident in AE1_OF-OF_, AE1_IF-IF_, and NBCe1. Both system- and state-specific differences in the magnitude of this enrichment are evident from the calculated DEI. Higher enrichment of CHOL and PIP2 lipids can be seen around the IF monomers of AE1_IF-IF_ than the OF monomers of AE1_OF-OF_. The average enrichment of PIP2 lipids around the OF homodimers of AE1_OF-OF_, NBCe1, and NDCBE decreases in the same order (the average DEI ranges from 5.1 in AE1 _OF-OF_ to 3.9 in NDCBE). The other negatively charged lipid, POPS, is depleted in all systems, likely due to competition with PIP2, which has a higher negative charge. POPS shows both state-specific (POPS depletion in AE1_IF-IF_ is lower than in the AE_OF-OF_ systems) and system-specific (POPS depletion decreases in the order AE1 < NBCe1 < NDCBE) trends. The enrichment of CHOL and PIP2 leads to depletion of the remaining lipid types in most studied systems, regardless of protein type and conformation state. A small system specific depletion can be seen in the distribution of POSM lipids, which are less depleted in NDCBE and negligibly depleted in NBCe1 (DEI ∼ 1.0). Small state-specific differences in depletion are observed in POPC distribution, where POPC lipids are slightly more depleted in the AE1_IF-IF_ systems compared to the AE1_OF-OF_ ones. Depletion of POPE lipids does not show system or state-specific trends.

**Figure 1.**
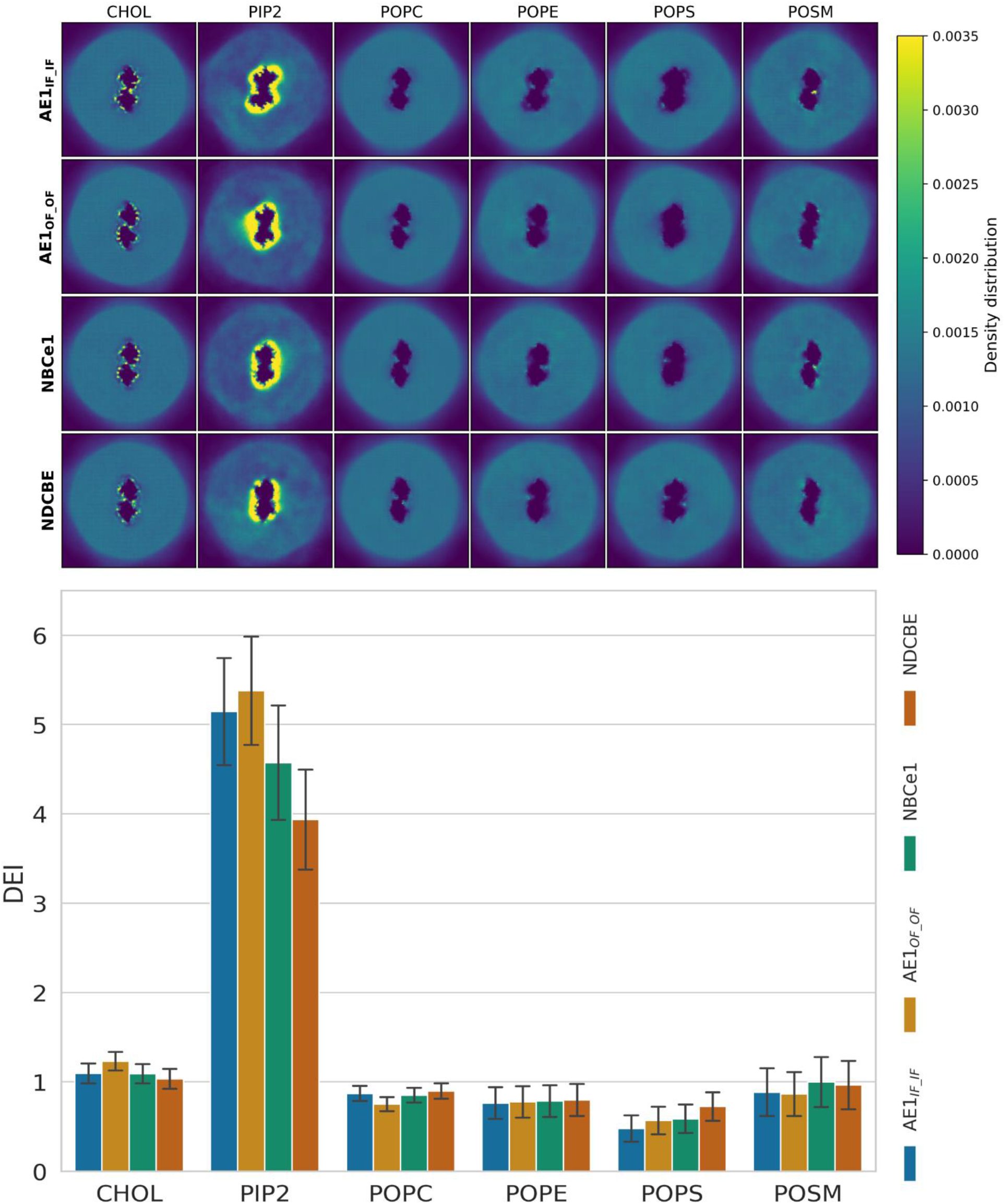
Density distribution and DEI of the four studied SLC4 members. **(top)** Density distribution plots and **(bottom)** calculated depletion and enrichment indices for different lipid types in the vicinity of the protein matrix of AE1_OF-OF_, AE1_IF-IF_, NBCe1, and NDCBE.

### Putative lipid coordination areas and binding sites in the protein matrix identified from contact analysis

#### a) Putative lipid coordination areas based on calculated occupancy and longest contact times

To assess in more detail the lipid organization around the protein matrix and to identify putative lipid coordination areas, occupancy and longest contact duration were calculated for all studied proteins at a 0.6 nm cutoff, using the ProLint server (43) as described in Ref.(42). For the purposes of the current work, occupancy (Figures S2-S7) is defined as percent of frames in which the headgroup of a lipid of a specific type is found within 0.6 nm of the protein, while longest contact duration (Figures S8-S13) maps the protein residues at which the headgroup of a lipid of a specific type spends the longest uninterrupted time (in µs) within a 0.6 nm cutoff. These two metrics capture frequent presence of lipids around specific residues and long-lived lipid-protein contacts and together can be used to map putative lipid coordination areas, where lipids of a specific lipid type are expected to be found more frequently and to bind for longer times. In addition, the occupancy plots (Figures S2-S7) provide evidence for convergence of the lipid-protein interactions in the analyzed final 20 µs of the simulations in the different replicas and between the monomers of the simulated homodimers (Chain A and Chain B in Figures S2-S13). Similar occupancy patterns can be seen both in comparison between different replicas and between the monomers in each dimer, especially in the cases of the lipid types with most prominent lipid-protein interactions (CHOL, PIP2, POPC). High occupancy values (∼100%) with a large number of protein residues in all studied proteins are observed for CHOL, PIP2, and POPC, showing that these lipids are in frequent contact with the proteins. The high occupancy value of POPC is not surprising, considering that this is the highest concentration lipid in the upper leaflet and one of the high concentration lipids in the lower leaflet; thus, frequent lipid-protein contacts with POPC would be expected. In the case of CHOL and PIP2, considering their lower overall concentration in the model HEK293 membrane used here, the high occupancy values indicate enrichment of the lipids around the protein, consistent with our DEI and density maps results in Figures 1 and S1. Despite its considerably higher concentration than PIP2, the other negatively charged lipid in our model membrane, POPS, shows lower occupancy values (∼50-80%) than the ones observed in the PIP2-protein interactions. Exceptions of this trend can be seen in some AE1_IF-IF_ and NDCBE replicas where POPS is in contact with a small number of protein residues in almost 100% of the trajectory steps. Such incidental high occupancy patterns may arise from entrapment of a POPS molecule in a monomer/replica during the simulation and are not necessarily a result of actual POPS binding sites in the protein matrix, especially considering the lipid density and enrichment-depletion data. POSM shows a ∼40% occupancy in most trajectories with the exception of the NBCe1 and AE1_IF-IF_ where higher occupancy values (∼50-100%) are consistently found in all hNBCe1 and AE1_IF-IF_ replicas/monomers, in line with the lipid density distribution plots in Figures 1 and S1. The longest contact trends mimic to some extent the occupancy trends. Generally, the longest-lived contacts are observed between CHOL and PIP2 lipids in all studied proteins. PIP2 contacts span the full 20 µs used for the analysis in most systems and replicas (except NDCBE where the contacts are slightly shorter), while CHOL contacts routinely exceed 10 µs (also evident upon trajectory inspection). POPC features long-lived protein/lipid contacts in some of the systems, where POPC molecules have bound to the dimerization interface (see below) and spend most of the simulation at that site. The remaining lipids (POPE, POPS, and POSM) have significantly briefer lipid-protein contacts, which are in line with previous simulations on various protein systems with these lipid types (42,44). Long-lived contacts of POPS and POSM are observed in the same monomers that feature high occupancy values for these lipids and may be a result of incidental lipid entrapment within the protein matrix.

A visual representation of occupancy and longest-contact duration can be done via heatmaps (Figures 2 and 3, respectively) which highlight the protein areas where these metrics are highest. Darker red color in Figures 2 and 3 indicate higher occupancy and longer contacts. Overlap of the intensely red areas from Figures 2 and 3 indicate potential binding sites for lipids of a specific lipid type. The occupancy plots (Figure 2) demonstrate that different lipids show preferences toward different parts of the protein although there is some expected overlap between POPC, POPE, and POSM and between POPS and PIP2, considering their chemical and structural similarities. Generally, CHOL is reliably found at the middle region of the lipid bilayer within a few well-defined clefts formed by the transmembrane helices (TMs) of the core domain (Figure 4) and, as previously reported in shorter CGMD simulations of AE1_OF-OF_ (18,19), at the dimerization interface. Frequent CHOL presence is also found in the area of TM13 of the gate domain in all occupancy maps. NBCe1, AE1_OF-OF_ and AE1_IF-IF_ show similar intense CHOL patterns in their CHOL longest contact plots (Figure 3), while NDCBE differs somewhat in intensity of the CHOL longest contacts in the core region and features a long-lived CHOL interaction in the area of TM13 at the protein dimerization interface. All three proteins show high occupancy and longest-contacts patterns for the PIP2 lipids, which are consistent with a stable PIP2 annulus at the intracellular side of the protein observed previously in shorter CGMD simulations in AE1_OF-OF_ (18,19). PIP2 lipids have highest occupancies and longest contacts at the dimerization interface or adjacent to it. PIP2 lipids are also involved in long contacts with the short H1 helix in all systems. POPS, which is the other anionic lipid in the simulation, competes with PIP2 for binding to the abundant positively charged protein residues in the lower leaflet (occupancy maps in Figure 2); however, PIP2 outcompetes it significantly. Nevertheless, POPS is still found to some extent in the same areas where PIP2 is observed in most systems. POPC is found predominantly around the protein at the upper leaflet (top of the membrane), where it competes with POPE and POSM and outcompetes them for contact with the protein surface (Figure 2) in line with its high concentration in the upper leaflet. Together with POPE and POSM, POPC features longer-lived contacts within the large cavity formed at the dimerization interface, indicative of possible lipid binding sites and participation of these lipids in the dimerization process.

**Figure 2.**
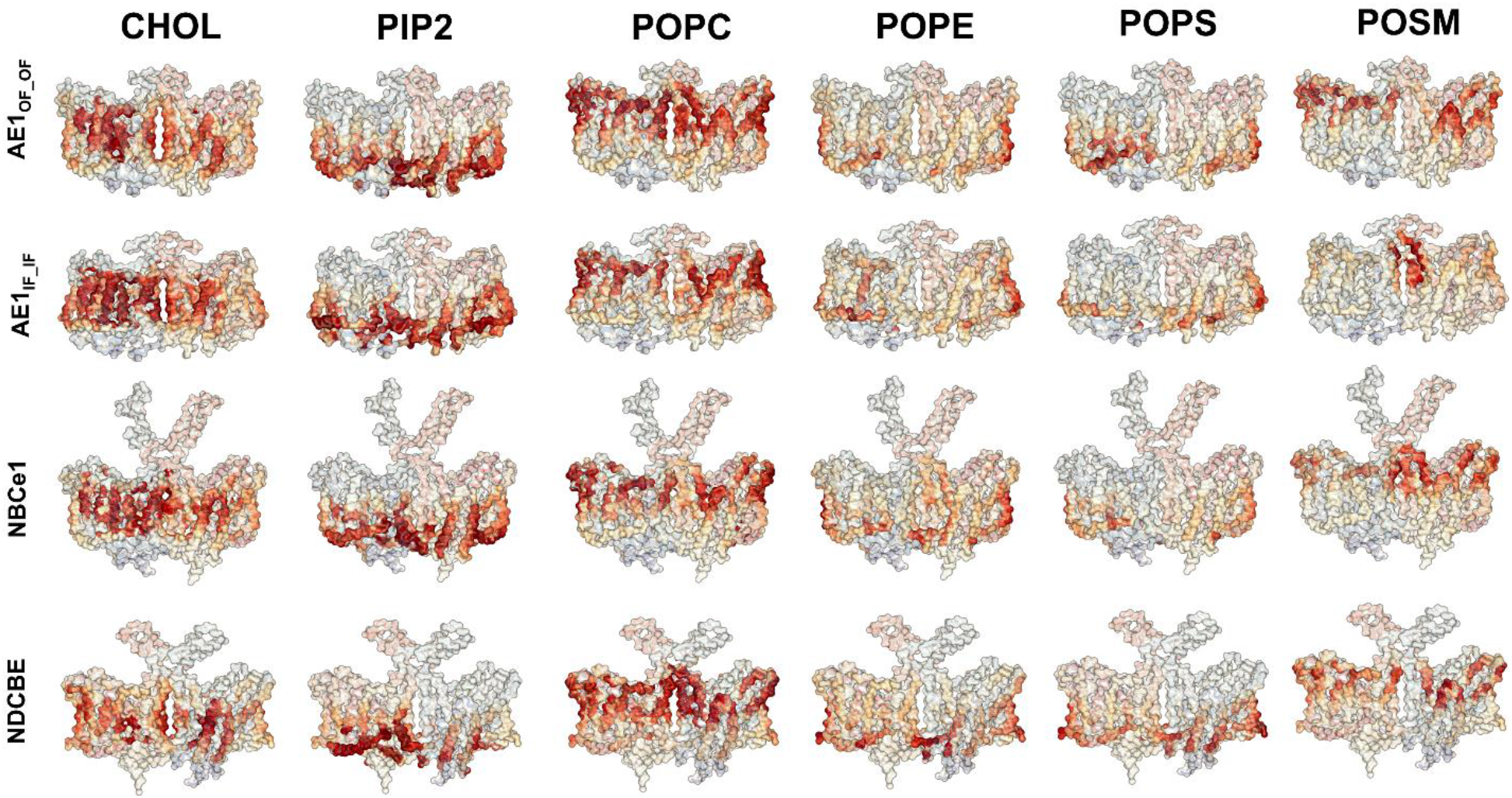
Lipid occupancy plots. Representative occupancy plots for different lipids in contact with the four studied SLC4 members, highlighting the preferred lipid-protein contact areas.

**Figure 3.**
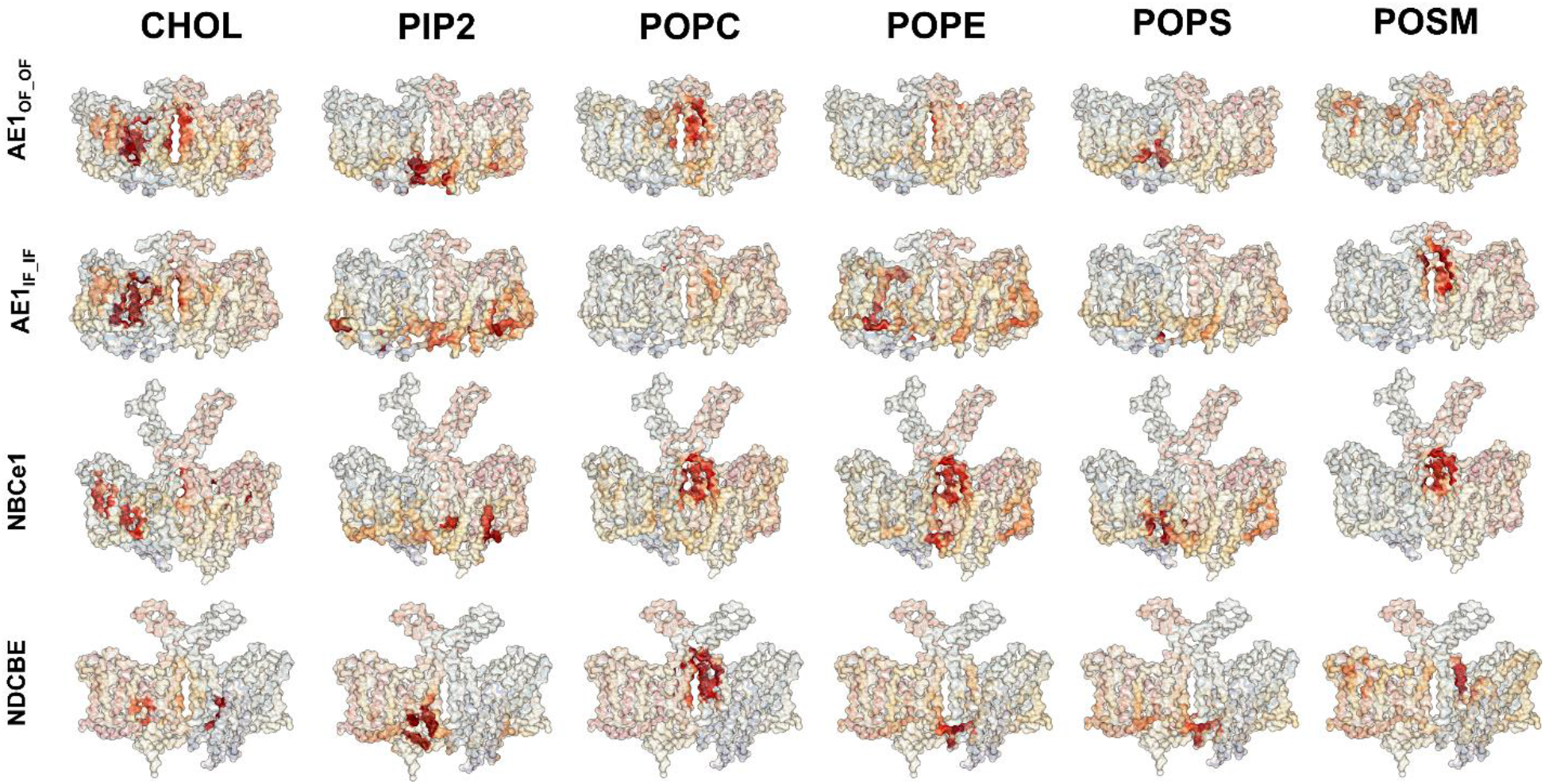
Lipid longest-contact plots. Representative longest-contact plots for different lipids in contact with the four studied SLC4 members, highlighting putative lipid binding areas.

**Figure 4.**
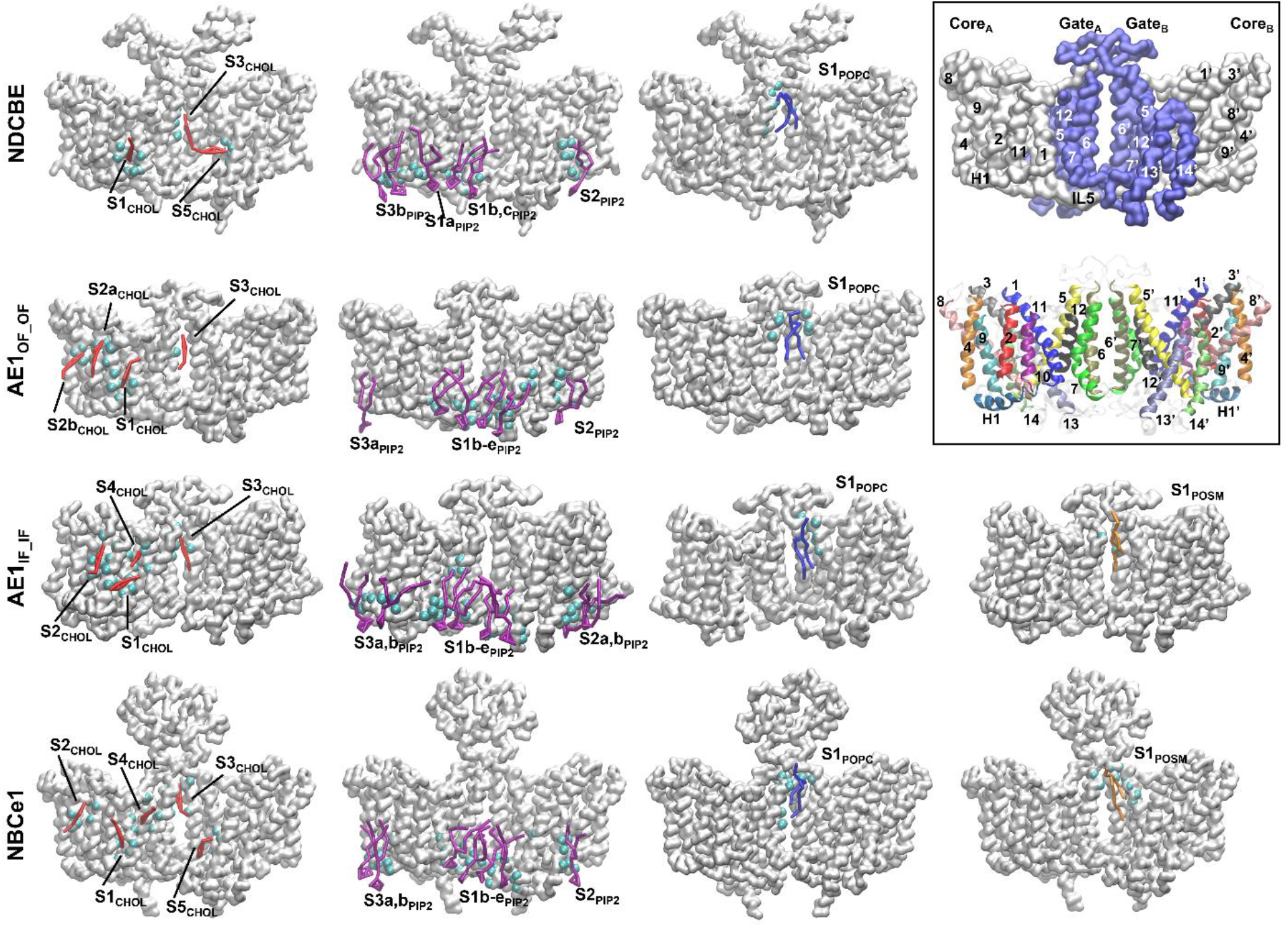
Putative lipid binding sites from coarse-grained MD trajectories. The lipids are color-coded as follows: CHOL in red, PIP2 in magenta, POPC in blue, POSM in orange. The position of the backbone atoms within 0.6 nm of the bound lipids are shown as cyan spheres. Since NDCBE and AE1_OF-OF_ did not show enrichment of the POSM around the protein matrix or long-lived protein-POSM interactions, we have not identified POSM binding sites in these two systems. **Excerpt:** Designation of the core and gate domains of monomers A and B of the SLC4 dimers and TM numbering, using AE1_OF-OF_ as an example. **(top)** Surface representation of AE1 divided into gate (blue) and core (white) domains; **(bottom)** Helical representation of AE1 with individually colored TMs.

### b) Putative binding sites for CHOL, PIP2, POPC, and POSM lipids based on shared contacts analysis

Figure 4 portrays lipid-protein contacts representing putative binding sites for the lipid types that show consistent preference for the SLC4 proteins based on density plots, DEI calculations, and contact analysis (e.g. CHOL, PIP2, POPC, and POSM) extracted from selected CGMD trajectories. The TM numbering and the core/gate designations of the monomers are shown overlayed with the three-dimensional structure of AE1_OF-OF_ as an excerpt in Figure 4. The position of the binding sites is determined from shared contact analysis, provided by the recently developed ProLint 2 GUI dashboard (45) and further validated with the occupancy and longest contact plots in Figures 2 and 3 and visual inspection of the MD trajectories. The lipids are shown in their coarse-grained representations and color-coded (CHOL in red, PIP2 in violet, POPC in blue, and POSM in orange). The position of the backbone carbon atoms within 0.6 nm of the lipids are indicated with cyan spheres in Figure 4 and are highlighted in the aligned protein sequences of human AE1_OF-OF_, bovine AE1_IF-IF_, human NBCe1, and rat NDCBE in the relevant lipid colors for comparison (Figure 5). In the OF systems CHOL molecules form frequent and long-lived contacts at residues from TMs 1, 2, 5, 6, 7, 9, 11, 13 and helices H3, H4 (Figures 3, 5, S8) of the core and gate regions, where the mutual orientation of the TMs results in hydrophobic clefts of suitable size and geometry for the CHOL molecules. CHOL binds also at the dimeric interface in all studied systems, in line with previous CGMD simulations of AE1_OF-OF_ (18,19). Overall, a total of five different putative CHOL sites can be identified in the protein matrix across the four studied protein systems, based on the longest contacts plots in Figure 3 and the shared contact analysis although the different systems show a different combination of these long-lived sites. However, these sites overlap with areas of high CHOL occupancy (i.e. frequent CHOL contacts), which exist in all four studies systems (Figure 2). Thus, they are consistently occupied by a CHOL molecule in the majority of the SLC4 trajectory steps even when the CHOL-protein contacts are brief and there is a high exchange of CHOL molecules at the sites. The core CHOL site labeled as S1_CHOL_ on Figure 4 is present in all studied systems in the cleft formed by H1 and TMs 1 and 11 (Figure 5). Long-lived CHOL-protein contacts at site S2_CHOL_ at the interface between TMs 2, 9 and 11 are observed in AE1_OF-OF_, AE1_IF-IF_, and NBCe1 although high frequency but shorter CHOL-protein contacts are also found in this area in NDCBE. Two frequently occurring positions for site S2_CHOL_ are shown (as S2a_CHOL_ and S2b_CHOL_) in AE1_OF-OF_ (Figure 4). A long-lived site S3_CHOL_ is consistently found at the dimerization interface of all proteins, between TMs 6 and 7 of the gate region. The dimerization interface opens a cavity-like structure which is permeated by multiple CHOL and lipid molecules throughout the CGMD simulations, thus only the highest occupancy/longest contact CHOL site is shown in Figure 4. Site S4_CHOL_ is present in AE1_IF-IF_ and NBCe1 in the area of TMs 1 and 7 and H3. In the AE1_IF-IF_ system, the rotation and vertical translocation of the core with respect to the gate domain leads to a deeper cleft at this area and a CHOL molecule can penetrate deeper between the AE1 core and gate domains along the short helix H3 (Figure 5) and bind for the full 20 µs of the analyzed trajectories. Thus, site S4_CHOL_ might have functional significance for the OF <-> IF conformational transition. A long-contact site S5_CHOL_ is identified only in NBCe1 and NDCBE in the area of TMs 5,13 and H4 from the gate region (Figure 4), although all systems feature frequent CHOL contacts in this area with similar CHOL orientation (Figure 2).

**Figure 5.**
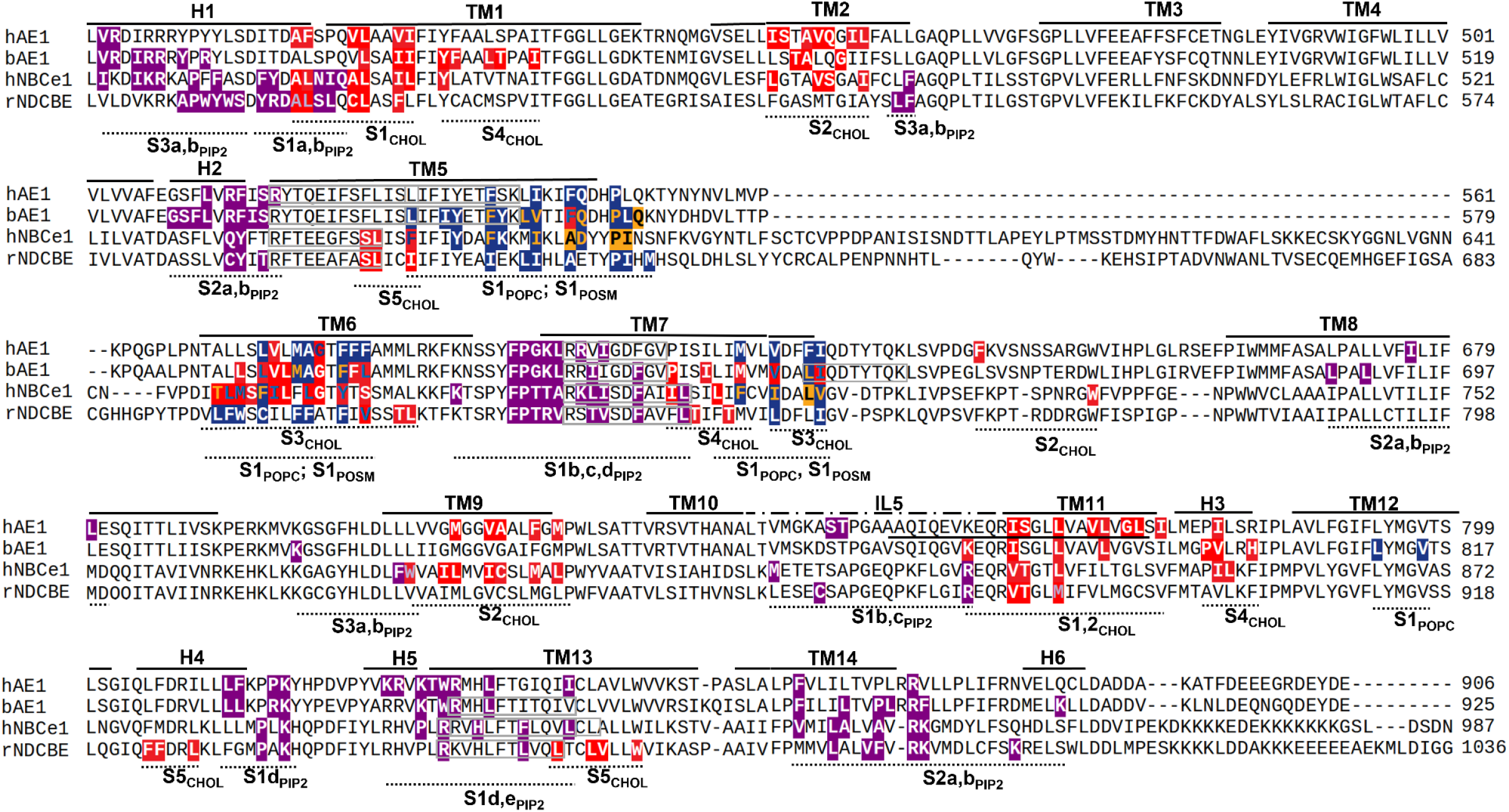
Sequence alignment of the four studied SLC4 systems. The identified putative lipid sites in **Figure 4** are highlighted in different colors, depending on the lipid type. The binding sites are formulated as backbone residues within 0.6 nm of the bound lipids (cyan sphere in **Figure 4**). The lipid binding sites are color-coded as in **Figure 4** (CHOL in red, PIP2 in magenta, POPC in blue, POSM in orange) and presence of more than one color at an amino acid residue signifies participation of this residue in more than one binding site. The CRAC/CARC domains in the areas of putative CHOL sites are displayed as grey boxes. See **Figure 4** for designation of the core/gate domains and TM numbering of the SLC4 monomers.

PIP2 forms multiple long-lived contacts at the intracellular side of all SLC4 proteins. We have identified 3 general areas (S1-3_PIP2_), where PIP2 lipids tend to accumulate and dwell (Figure 4) during the MD trajectories. Site S1_PIP2_ is a wide strip of residues at the dimerization interface and includes residues from TMs 1, 7, 13 and helices H1, H4 and H5, which feature multiple basic residues available for contact with negatively charged lipid heads (Figure 5). This area can be split further into 5 smaller binding sites (S1a-e_PIP2_, Figure 4) and is usually occupied by 2-4 PIP2 molecules during the CGMD simulations. Sites S1b-e_PIP2_ are present in all studies systems (Figure 5). In NDCBE and NBCe1 the PIP2 molecules form longer contacts also at site S1a_PIP2_, while in the remaining two systems, AE1_OF-OF_ and AE1_IF-IF_, site S1a_PIP2_ forms more transient contacts. Instead, sites S1d,e_PIP2_ are more frequently occupied and for longer times in these systems. Residues from the intracellular loop 5 (IL5), between TMs 10 and 11 are also involved with PIP2 interactions mostly at sites S1b,c_PIP2_. In the OF states, IL5 forms a β-hairpin structure, tucked under the protein where it can interact with the PIP2 lipid heads at the core-gate interface. In AE1_IF-IF_ due to the lateral motion of H1, the downward motion of the core and TM10, and the elongation of TM11, during which some IL5 residues are incorporated into the α-helical shape of TM11 (27), the points of contact between IL5 and PIP2 decrease and residues from the beginning of IL5 become available for rare and transient PIP2-protein contacts (Figures 4-6). A second binding area, S2_PIP2_, positioned between TMs 8, 14 and helices H2, H6, is found in all four studied systems. This site is consistently involved in a long-lived contact with at least one PIP2 molecule in most monomers but in AE1_IF-IF_ due to the rotation and vertical displacement of the core, a second long-lived PIP2-protein contact can be observed as well (Figure 4, 6). The last PIP2 binding area, site S3_PIP2_ is found at H1 in all systems and is especially prominent in AE1_IF-IF_ where it becomes available for long-lived contacts as H1 moves laterally from underneath the protein (27) (Figures 4, 5). Two long-lived PIP2 binding sites, S3a,b_PIP2_, can be distinguished in this case (Figure 4).

**Figure 6.**
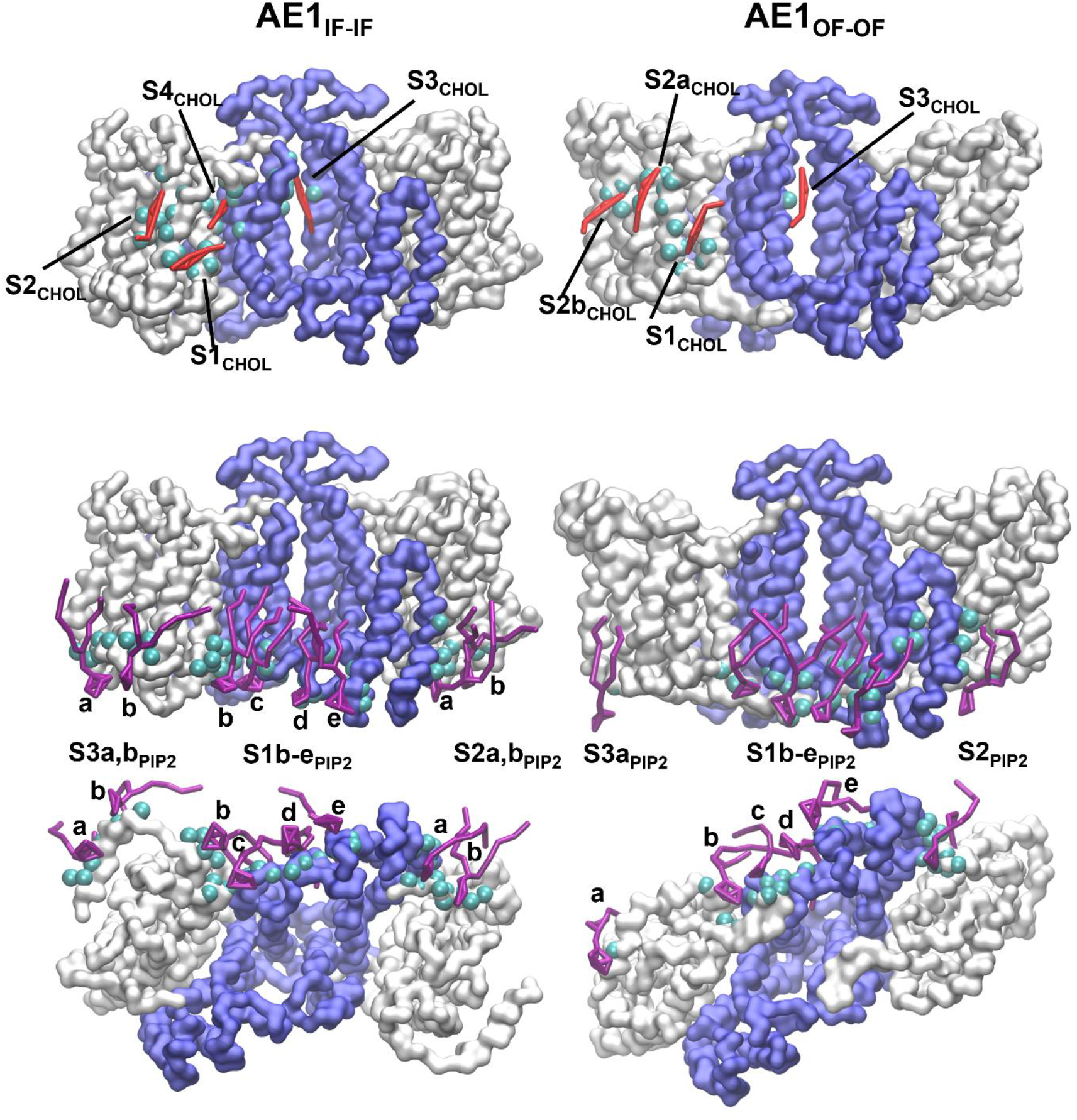
Lipid participation in the conformational transition of AE1. Potential participation of CHOL and PIP2 in the OF <-> IF transition in AE1. **(top)** Side view of CHOL binding sites in AE1_IF-IF_ and AE1_OF-OF_. **(bottom)** Side and bottom view of PIP2 binding sites in AE1_IF-IF_ and AE1_OF-OF_. Color-coding: core domain as white surface, gate domain as blue surface, CHOL molecules as red sticks, PIP2 molecules as magenta sticks, backbone atoms within 0.6 nm of the bound lipids as cyan spheres. See **Figure 4** for designation of the core/gate domains and TM numbering of the SLC4 monomers.

POPS interacts with the same binding areas as PIP2 (Figure 2 and 3); however, considering its depletion around the protein matrix (Figure 1) due to the competition with the highly charged PIP2, this leads to rare occurrence of long-lived contacts. Nevertheless, such contacts can be identified in the areas of sites S1-3_PIP2_ in AE1_IF-IF_ and NDCBE (Figure 2).

The binding sites of the remaining two lipids which form long-lived contacts of potential significance (considering also density distribution and DEI, Figure 1) with the SLC4 proteins, POPC and POSM, can both be found at the dimeric interface (sites S1_POPC_ and S1_POSM_, Figure 4) in the areas of TMs 5, 6, and 7 of the gate region, where they overlap also with sites S3,4_CHOL_ (Figure 5). The POPE lipids, which are depleted around the protein matrix and do not feature long contacts with any of the studied proteins, generally tend to interact with similar residues as POPC and POSM and can also be found in the areas of sites S1_POPC_ and S1_POSM_ (Figure 2).

## Discussion

### Similarities and differences in lipid-SLC4 interactions from the identified lipid/CHOL binding sites

Occupancy plots (Figure 2) show similar occupancy trends for all studied lipid types in all three proteins. However, the longest contact plots (Figure 3) display a more varied picture of the lipid-protein contacts and the extracted binding sites based on these plots and shared contacts analysis, together with the system-specific trends in lipid densities and DEI, point at subtle differences between the different members of the family. The functional relevance of these differences and whether they can be related to lipid regulation of SLC4 function is yet unclear. Nevertheless, the newly identified lipid binding sites provide a basis for future functional mutagenesis studies which can shed more light on the potential role of lipids in the protein targeting to the membrane, oligomerization, and transport mechanism. Significant system-specific trends in contact frequency, longest contact duration, density distribution, and DEI are observed for CHOL, PIP2, and POSM lipids with NDCBE showing more distinct occupancy and longest-contact patterns from the remaining studied systems (AE1 and NBCe1). AE1 and NBCe1 show generally similar protein-lipid interaction trends, although with slightly varying magnitude (smaller enrichment of PIP2 and CHOL around NBCe1, more prominent presence of POSM at the lipid interface in NBCe1 than in OF AE1). NDCBE is expressed and functional in neurons (7,26), while AE1 and NBCe1, are found in the distal and proximal tubule of the nephrons where they participate in urine acidification (5,7,9,20). AE1 is also heavily expressed in erythrocytes (9). Thus, the different cellular membrane composition in neurons, the distal and proximal tubule of nephrons, and the erythrocytes could have been a driving force behind the development of slightly different lipid-protein fingerprints between these three members of the family.

The membrane compositions of model erythrocyte, neuron, average cell, and our HEK293 model membrane are shown in Table S1. The experimental lipidomics data for the basolateral membrane of the proximal tubule is included in the table as well. There are significant differences in the lipid composition between the different cell types. Most notably, the erythrocyte membrane has higher SM, PS, and CHOL content than the rest. The neuronal membrane features a higher concentration of PI lipids than the remaining cells even though NDCBE demonstrates the smallest PIP2 enrichment of all systems. The basolateral membrane of the proximal tubule has the largest amount of PE lipids, although these lipids did not show any system or state-specific trends in our study. Nevertheless, cross-referencing the membrane compositions in Table S1 and the observed system-specific trends in our simulations do not provide unambiguous answers about the mechanism of lipid regulation of SLC4 function and if such regulation exists. More detailed studies of the overall protein-lipid effects in the SLC4 family and of the role of the putative binding sites identified in this work would be required for correlation of the provided membrane compositions in different cell types, the behavior of the lipids in the simulations reported here, and the overall SLC4 function.

### Experimental evidence for the proposed binding sites

A number of mutations which impact the AE1 and NBCe1 function have been discovered at the protein/lipid interface, in the areas of some of the lipid binding sites identified here. A list of these mutations and their relevant lipid binding sites are presented in Tables S2 and S3. Although the exact mechanism of the effect of these mutations on the protein function is rarely known, their location in the vicinity of the putative lipid binding sites demonstrates the functional relevance of these protein areas.

The most prominent SLC4 interactions are with the two lipid types that show consistent and significant enrichment around the protein matrix (Figures 1, S1) and/or high occupancy and longest-contacts trends (Figures 2, 3, S2, S3, S8, S9) - PIP2 lipids, forming a stable lipid annulus at the intracellular side of the proteins, and CHOL, spanning the whole width of the lipid membrane and accumulating at several binding sites in the core and gate regions of the proteins. We identify a total of 5 CHOL binding sites in the core and gate areas of AE1_OF-OF_, NBCe1, and NDCBE (Figure 4). Simultaneous binding of two (NDCBE) to four (AE1, NBCe1) CHOL molecules per monomer is routinely observed during the CGMD simulations. Binding of three CHOL molecules per monomer to the core region of AE1_OF-OF_ has been demonstrated also in a recent cryoEM structure resolved in lipids from Sf9 insect cell lines (46). In addition, single CHOL binding sites in the junction area of H5 and TM13 have been reported in recent cryoEM structures of AE1 and AE2 (47,48). The experimentally determined CHOL sites are in the general areas of sites S1_CHOL_, S4_CHOL_, S2_CHOL_, and S5_CHOL_ (Figure 4), although their exact position and the orientation of the CHOL molecule within them are not perfectly reproduced due to the coarse-grained nature of our simulations.

CRAC/CARC domains have been proposed in the literature on lipid-protein contacts as potential amino acid motifs selective for CHOL, although they suffer from high rates of both false positives and false negatives (49). The sequences of human AE1, bovine AE1, human NBCe1, and rat NDBCE used in this study possess several amino acid motifs, consistent with CRAC/CARC domain sequences (Table S4), as determined by the ScanProsite tool (50). However, most of these motifs were found in areas that do not feature CHOL-protein interactions in our simulations. The CRAC/CARC-like motifs found in the vicinity of putative CHOL binding sites are highlighted with a grey box in Figure 5, along with their corresponding CHOL sites. Conserved CRAC/CARC-like motifs associated with CHOL binding are found in all four systems at the gate domain, in the areas of TMs 5, 7 and 13, in the vicinity of site S4_CHOL_, which may have a functionally significant role in the OF <-> IF transition (see below) and site S5_CHOL_, which is one of the CHOL sites resolved in several different cryoEM structures of AE1 and AE2 (46-48). CRAC/CARC-like motifs also appear at the dimerization interface between TMs 5 and 7, in the vicinity of sites S1_POPC_ and S1_POSM_, which are in proximity to site S3_CHOL_. In line with this, the dimerization interface experiences frequent contacts with CHOL molecules, as seen in Figure 2 and in previous simulations (18,19). However, multiple CHOL binding sites are identified in areas devoid of CRAC/CARC motifs, with relatively low sequence identity between the four studied systems, that nevertheless host a variety of branched hydrophobic, aromatic and polar residues and provide an appropriate three-dimensional geometric fit with the CHOL molecules, as previously observed in GPCR simulations (42). Among these sites is site S1 which was identified in all four systems, regardless of the absence of a CRAC/CARC motif in this area. This suggests that the CHOL-protein interactions in the SLC4 family are dictated at least partially by mutual alignment of the TMs, which open suitably-sized clefts at the lipid-protein interface and which are reproduced in all SLC4 proteins and, likely, other 7+7 fold proteins.

CryoEM structures of AE1_OF-OF_ with a single PIP2 molecule bound in the area of site S1b,c_PIP2_ identified from our simulations was also recently reported (47). Multiple phospholipid types (POPC, POPE, POSM) and CHOL compete for binding at the dimerization interface of the gate domains (Figure 4) as determined here and in previous simulations on AE1_OF-OF_ dimers (18,19). Phospholipids binding in the area of sites S3_CHOL_, S1_POPC_, and S1_POSM_ have been reported in cryoEM structures of AE1 (46,47) in line with our findings.

### Previous experimental data on SLC4-lipid interactions in the context of our computational findings

Most previous work on lipid-protein/CHOL interactions in the SLC4 proteins involved the AE1 and NBCe1 members of the family. Our simulations demonstrate that CHOL is enriched in most studied SLC4 systems (except NDCBE, where this enrichment is negligible) and is involved in both frequent and long-lasting interactions with the protein matrix (Figures 2, 3) leading to identification of several putative CHOL binding sites (Figure 4). In line with this, specific CHOL interactions with AE1 have been demonstrated with ESR and fluorescence photo-bleaching recovery experiments by “immobilization” and decreased diffusion of CHOL in AE1-containing erythrocyte ghost membranes (17) and liposomes (13). Surface pressure measurements in monolayers of various lipid mixtures suggest that CHOL forms strong interactions with AE1, which may be modulated by other lipids in the mixture (10,11). In our simulations, the CHOL binding areas overlap with the binding areas of other lipids, such as PIP2, POPC, and POSM (Figure 5). Competition for these binding areas may give rise to the observed lipid modulation of CHOL interactions. Existence of a CHOL binding site in AE1 which inhibits anion transport has also been proposed (11). A possible candidate for this site is S4_CHOL_, which may play a role during the OF <-> IF transition (see below). Similar effects of low PIP2 diffusion due to PIP2-AE1 interactions have been detected with pyrene fluorescence experiments in DOPC vesicles with varying PI, PIP and PIP2 content (14) and are in agreement with the high enrichment of PIP2 around all studied SLC4 systems and the consistently long PIP2-protein contacts which span the full 20 µs of the analyzed trajectories in the areas of the identified PIP2 binding sites (Figures 2, 3, S9). Patch-clamp current measurements in HeLa cells and Xenopus oocytes have demonstrated that different isoforms of NBCe1 (A, B, and C) are stimulated to different degree by cytosolic PIP2, hinting at possible PIP2 regulation of NBCe1 (21-23). A putative PIP2 binding site at the positively charged residues 37-65 of the N_t_ terminus of NBCe1-B (part of the autoinhibitory domain of NBCe1-B) was also identified with functional mutagenesis experiments (23). Unfortunately, this highly flexible area of NBCe1 has never been resolved and is missing from our models. However, involvement of the PIP2 annulus in the PIP2 regulation would not be surprising considering that the identified S1_PIP2_ binding sites are in proximity to the IF cavity opening (27) which may be blocked by the PIP2 bound N_t_ during auto-inhibition. Fluorescence digital imaging microscopy on formation of lipid domains influenced by hAE1 and spectrin, demonstrates better correlation between AE1 and PC domains than between AE1 and PS domains (i.e. stronger AE1 interactions with PC than PS) (15). Both POPC and POPS are depleted around the protein in our simulations (Figure 1), displaced by CHOL and PIP2, respectively. POPS is found in the same areas at the intracellular side of the SLC4 proteins where PIP2 tends to accumulate, attracted by a multitude of positively charged residues and is expected to form strong lipid-protein interactions in these areas in the absence of PIP2, as seen in previous simulations of AE1 in POPC/POPS lipids (19). The stronger interactions of AE1 with PC then may be due to the long-contact lipid binding at site S1_POPC_ at the dimerization interface, which is observed in most of the simulated SLC4 replicas (Figures 2-4) and by the more uniform contact coverage of the protein matrix by the POPC molecules (Figure 2). Nevertheless, the binding patterns observed in our work, where the lipids are homogenously spread, have the same type of lipid tails, and do not form isolated lipid domains, may not be directly comparable to these results. Previous computational modelling of AE1 in phase-separated membranes have shown tendencies of AE1 to sequester differently into low and high-order lipid domains (51); thus inclusion of phase-separation in the model membranes may be important for a more realistic description of the lipid-protein interactions in the SLC4 family. Dependence of hAE1 ^35^SO_42-_/SO_42-_ exchange on the lipid composition of vesicles shows that AE1 transport is insensitive to replacement of PC by PE, while SM and PS inhibit AE1 transport at higher concentrations. Rigidification of the membrane due to saturated lipid tails and formation of H-bonds between the lipids and the protein, electrostatic repulsion between anions and negative lipid heads, and lipid-modulated change in the aggregation of hAE1 into dimers and tetramers have been suggested as possible explanations of these effects (16). In our simulations, POPE is depleted in all systems and does not show system or state-specific trends. Its binding patterns are generally similar to those of POPC (Figure 2). Thus, the two lipids may be undistinguishable when it comes to impact on anion transport. This presumed impact may be solely determined by their relative concentrations. The binding sites of POPS (and PIP2) are at the intracellular side, at the dimerization interface, which has been previously identified as an anion reservoir, relevant to anion permeation into the IF state of AE1 (27). Presence of negatively charged lipids in this area may then indeed interfere with anion accumulation in the reservoir and successive entry in the IF cavity. POSM is less depleted than POPS around all systems and in the case of NBCe1, this depletion is negligible. POSM also shows high density at the dimerization interfaces of NBCe1 and AE1_IF-IF,_ which corresponds to the identified site S1_POSM_ in Figure 4. Thus, POSM and other phospholipids may be involved in modulation of the dimerization processes in transporters of the 7+7 architecture as previously suggested (52-54) and observed in multiple oligomers of secondary transporters (2,54,55). The effect of oligomerization on SLC4 transport is not clear. Structural and electrophysiological evidence shows that the individual monomers in SLC4 (and likely other proteins with the same 7+7 architecture) can function independently from one another (27,56). Oligomerization in secondary transporters, however, likely increases the efficiency of transport, as demonstrated previously in LeuT-fold proteins (57). Thus, phospholipid (e.g. POPC, POSM, POPE) binding at the dimerization interface may indeed modulate transport indirectly, by modulating oligomerization.

### Potential lipid role in the OF <-> IF conformational transition

Figure 6 compares CHOL (side view) and PIP2 (side and bottom view) binding in AE_OF-OF_ and AE1_IF-IF_ dimers. The OF <-> IF transition can be regarded as a rigid elevator motion of the core domain (white surface in Figure 6) with respect to the immobile gate domain (blue surface in Figure 6), which serves as the dimerization scaffold in the SLC4 dimers (27). A notable difference between the IF and OF states of AE1 is the existence of a long-lived CHOL site, S4_CHOL_, in the IF state. The CHOL molecule in S4_CHOL_ has penetrated the deep cleft formed between the core and gate domains and remains there for the full 20 µs of the analyzed MD trajectory in most of the AE1_IF-IF_ monomers. Frequent contacts consistent with site S4_CHOL_ appear also in the OF states of the remaining SLC4 systems, including NBCe1, where the CHOL-protein contacts in this area are also among the longest-lived ones. However, in the OF states, the S4_CHOL_-bound CHOL molecule does not permeate deep along helix H3 and occasionally dissociates and exchanges with other CHOL molecules in the span of the 20 µs trajectories. Considering the position of site S4_CHOL_ between the core and gate domains and the behavior of the CHOL bound at this site, it is possible that site S4_CHOL_ is the previously proposed CHOL inhibitory site (11), which in this case would inhibit transport by locking AE1 in its IF conformation. A possible mechanistic role of this site is the CHOL-mediated stabilization of the IF state which is of higher energy (and likely less stable and shorter lived) than the OF state, as determined by previous coarse-grained metadynamics simulations (27). In erythrocytes and erythrocyte ghost membranes, which feature higher concentration of CHOL than the model HEK293 membrane used in this study, AE1 has been suggested to assume predominantly the IF state (58-60).

Other lipid sites with possible involvement in the OF <-> IF transition are sites S1b,c_PIP2_ at the interface between the core and the gate and in the area of TM11 and the intracellular loop IL5, connecting TMs 10 and 11. This region is crucial for the OF <-> IF transition in AE1 and oscillates between an α-helical (in the IF state where TM11 is elongated) and a β-hairpin meta-stable state (when TM11 is partly unfolded) (27). The PIP2 lipids in sites S1b,c_PIP2_ thus interact with different portions of the intracellular loop as the monomers switch between the OF and the IF state. The conformational changes in TM11 and the intracellular loop are accompanied also by motion of the short helix H1 which shields the IL5 residues incorporated in the elongated portion of TM11 from the surrounding PIP2 lipids. Instead, the S3a,b_PIP2_ sites become more prominent and long-lived once H1 has shifted laterally and has uncovered several positively charged residues which were previously shielded by TM11 and TM2 of the core domain. Thus, PIP2-protein interactions in this area might have mechanistic significance for the concerted motion of TM10-IL5-TM11-H1 during the conformational transition, including involvement into PIP2 regulated auto-inhibition in NBCe1 (see above) once the IF cavity opens toward the cytoplasm.

## Conclusions

In this work we performed computational assessment of the lipid behaviour around the protein matrix and identified putative binding sites for different lipid types in three members of the SLC4 family, AE1 (IF and OF states), NBCe1 (OF state), and NDCBE (OF state). The most prominent protein-lipid interactions in all studied proteins are with CHOL and PIP2 lipids which show increased density distribution and are enriched around the protein matrix. System- and state-specific trends are observed in the density distribution, DEI, contact frequency, longest contact duration, and shared contact analysis for CHOL, PIP2, POPC, POPS, and POSM. NDCBE shows more distinct patterns of interaction with CHOL and PIP2 than AE1 and NBCe1. Five CHOL binding sites are identified in suitably organized clefts between TMs of the core domain and in the large cavity at the dimerization interface. PIP2 forms an annulus at the intracellular side of all studied proteins, which can be divided into three multi-lipid binding sites. The distinct contact patterns of CHOL and PIP2 in the IF and OF states of AE1 indicate that these two lipids might have mechanistic significance for the IF <-> OF transition in the SLC4 family. A single long-lived overlapping binding site is identified for CHOL, POPC and POSM at the extracellular side of the dimerization interface implying a potential role of these lipids in the SLC4 oligomerization. Our results are in good agreement with recently identified CHOL, PIP2 and phospholipid sites in AE1 and AE2 cryoEM structures and with available prior experiments on lipid effect on AE1 and NBCe1 function.

## Author contributions

H.R.Z., D.P.T., A.P, and I.K designed the study. H.R.Z. performed the coarse-grained simulations. H.R.Z., D.R.E., and B.I.S. performed the analysis of the simulated trajectories. D.R.E. and B.I.S. developed and modified the ProLint server for the purpose of the present trajectory analysis and deposited the SLC4 trajectories and results in the online ProLint database. H.R.Z. wrote the first version of the manuscript. All authors contributed to the final version of the manuscript.

## Acknowledgments

I.K. is supported by the NIH grant R01DK077162, the Factor Family Foundation Chair in Nephrology, Smidt Family Foundation, Paula Block Charitable Foundation, and the Kleeman Foundation. We acknowledge the Department of Medicine for support of A.P. Work in the D.P.T lab was supported by the NIH (R01 DK077162) and by Canadian Institutes of Health Research (CIHR, grant PJT-180245) and the Canada Research Chairs programs. MD simulations were performed on the Digital Research Alliance of Canada (previously known as Compute Canada) clusters under Research Allocation Award to D.P.T. H.Z. would like to thank Haydee Mesa-Galloso for providing the scripts for DEI calculations.

## Supporting Material

The supporting material includes density plots and protein-lipid contact figures for all replicas, tables with lipid compositions of different cellular and model membranes, known pathological mutations in AE1 and NBCe1, and relevant CRAC/CARC domains in the four studied proteins. The analyzed 20 µs trajectories for all replicas have been deposited in the ProLint results database at https://www.prolint.ca/explore/slc4 and are available for interactive viewing of the results reported here with the ProLint visualization tools.

## Supporting Citations

References (61-68) appear in the Supporting material

