## Supporting Material for "Coarse-grained molecular dynamics simulations of lipid-protein interactions in SLC4 proteins"

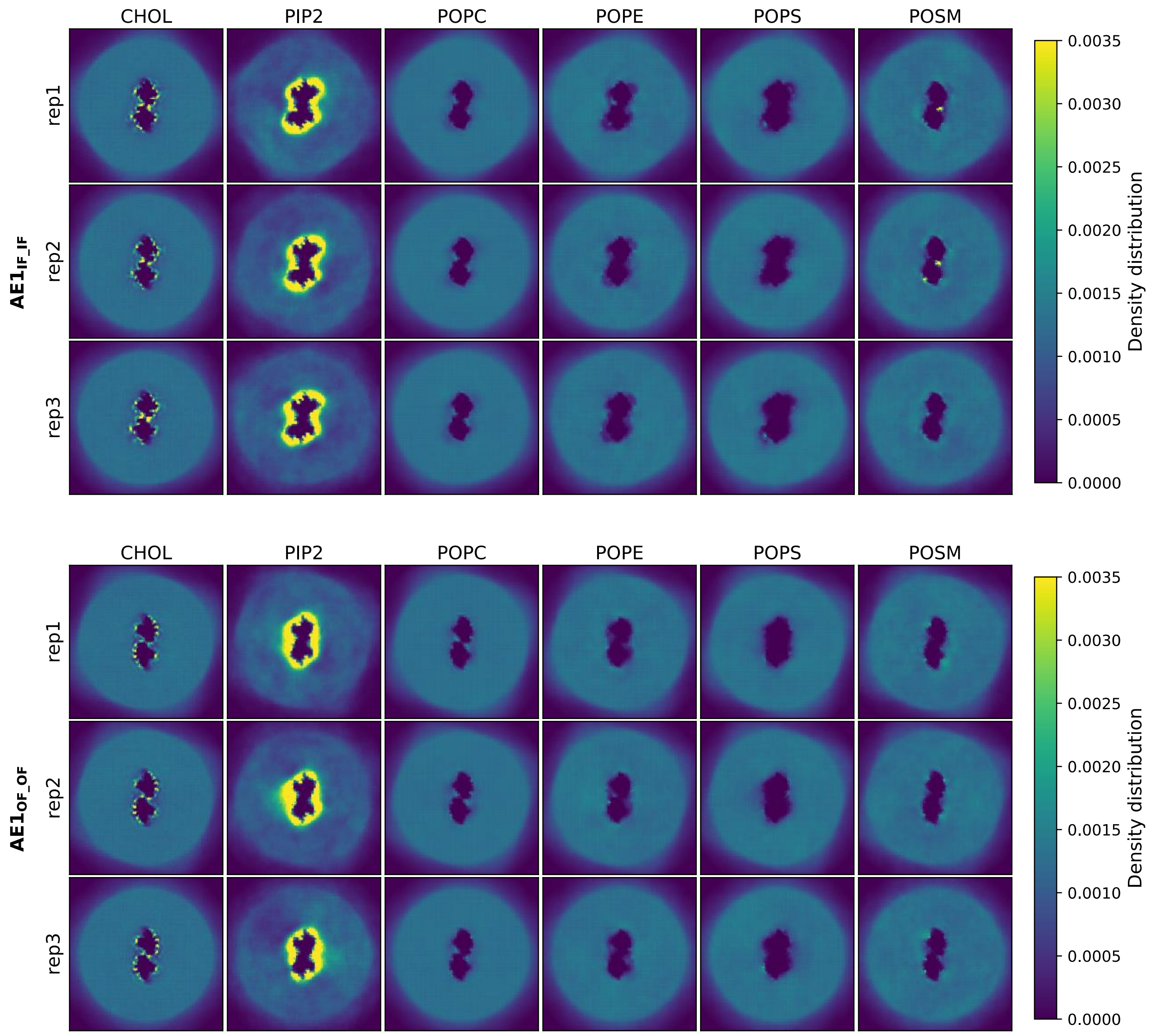


**Figure S1** Density distribution plots for different lipid types in the vicinity of the four studied SLC4 members. Comparison of all simulated replicas.

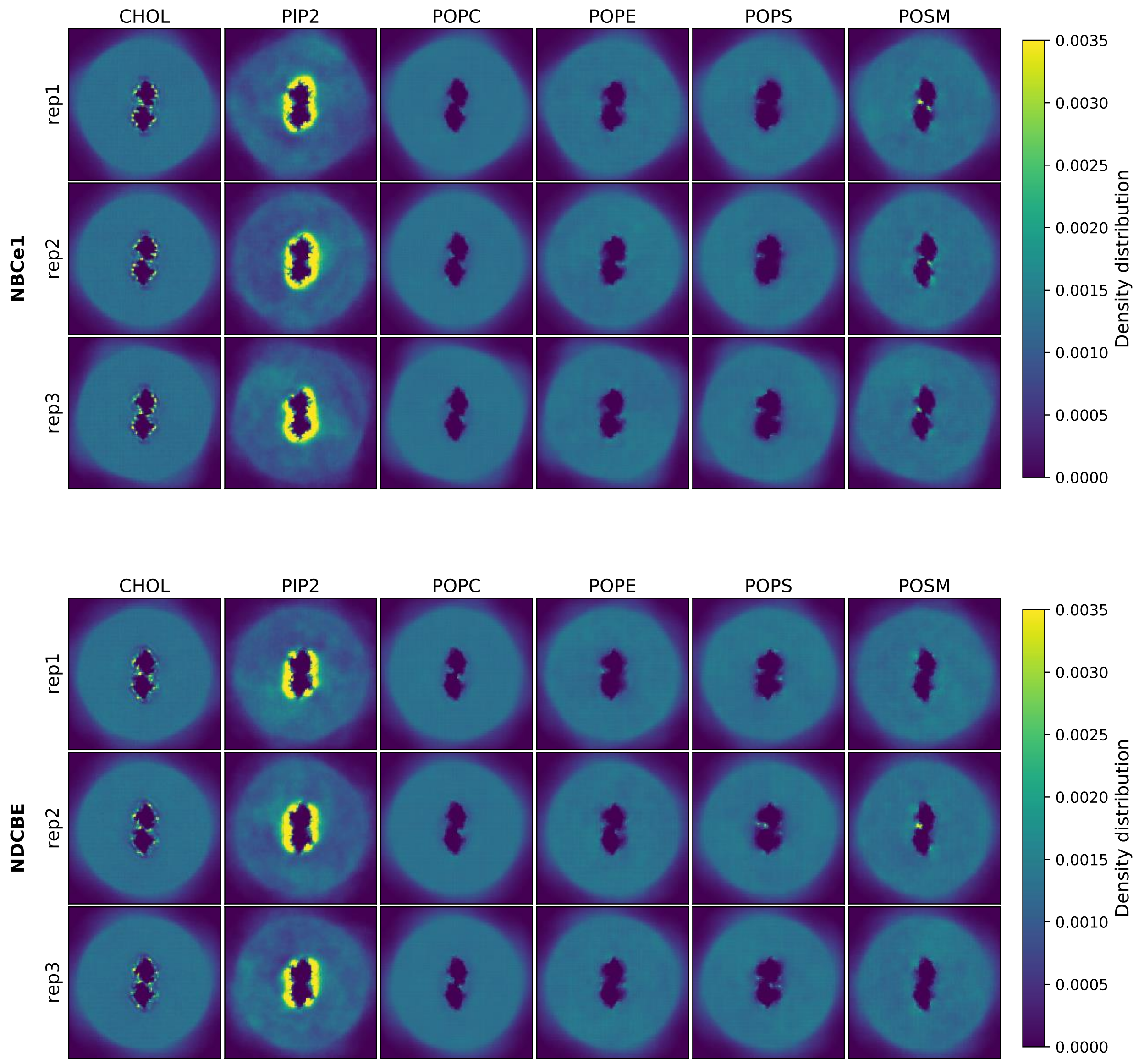


**Figure S1 (continued)** Density distribution plots for different lipid types in the vicinity of the four studied SLC4 members. Comparison of all simulated replicas.

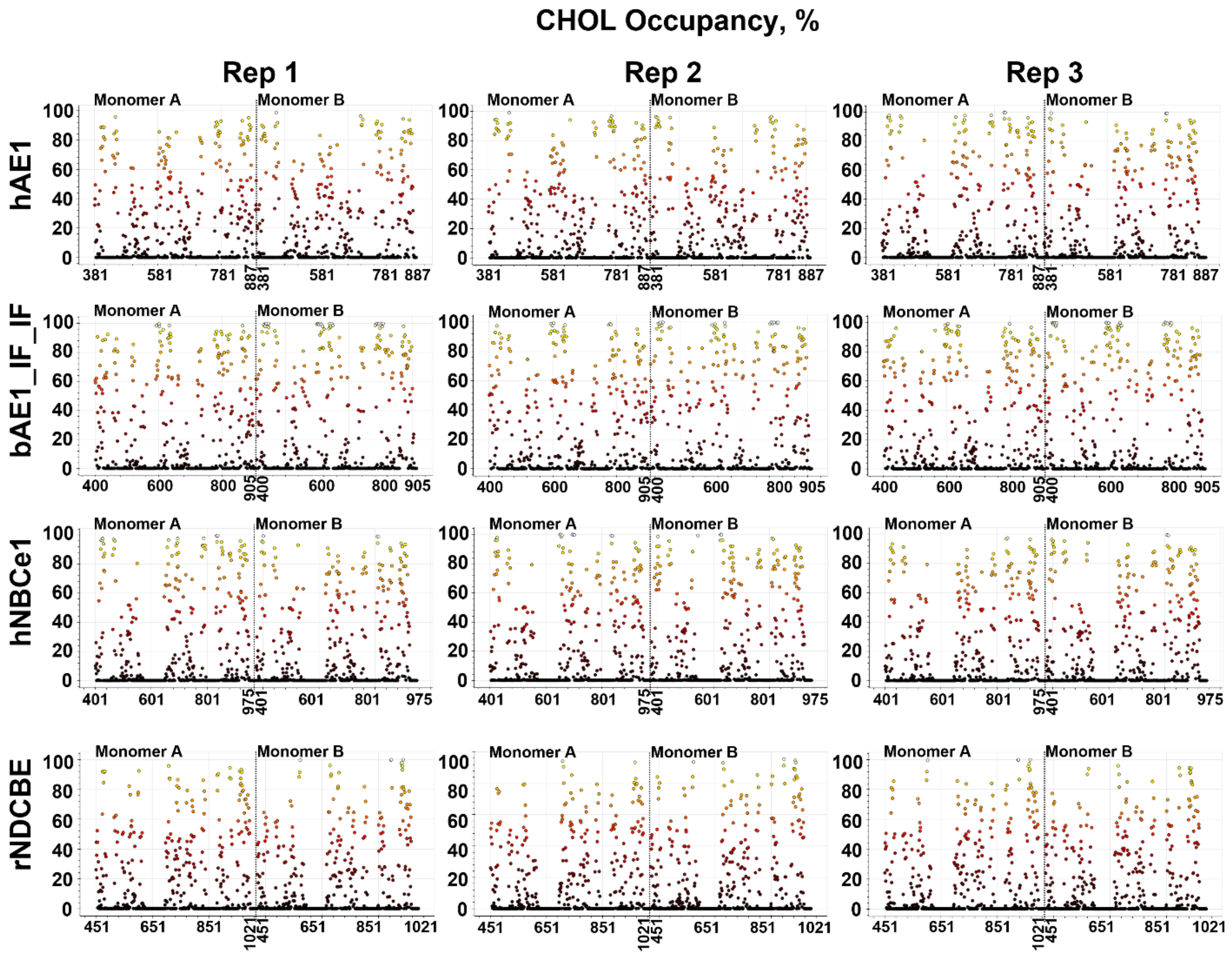


**Figure S2** Occupancy plots (in %) for CHOL-protein contacts in all simulated replicas (three replicas Rep1 – Rep3 per protein dimer). The end of the plot for the first monomer in the dimer (Monomer A) and the beginning of the plot for the second monomer of the dimer (Monomer B) is signified by a dashed line and the provided numbering of the amino acid residues along the x-axes of the plots.

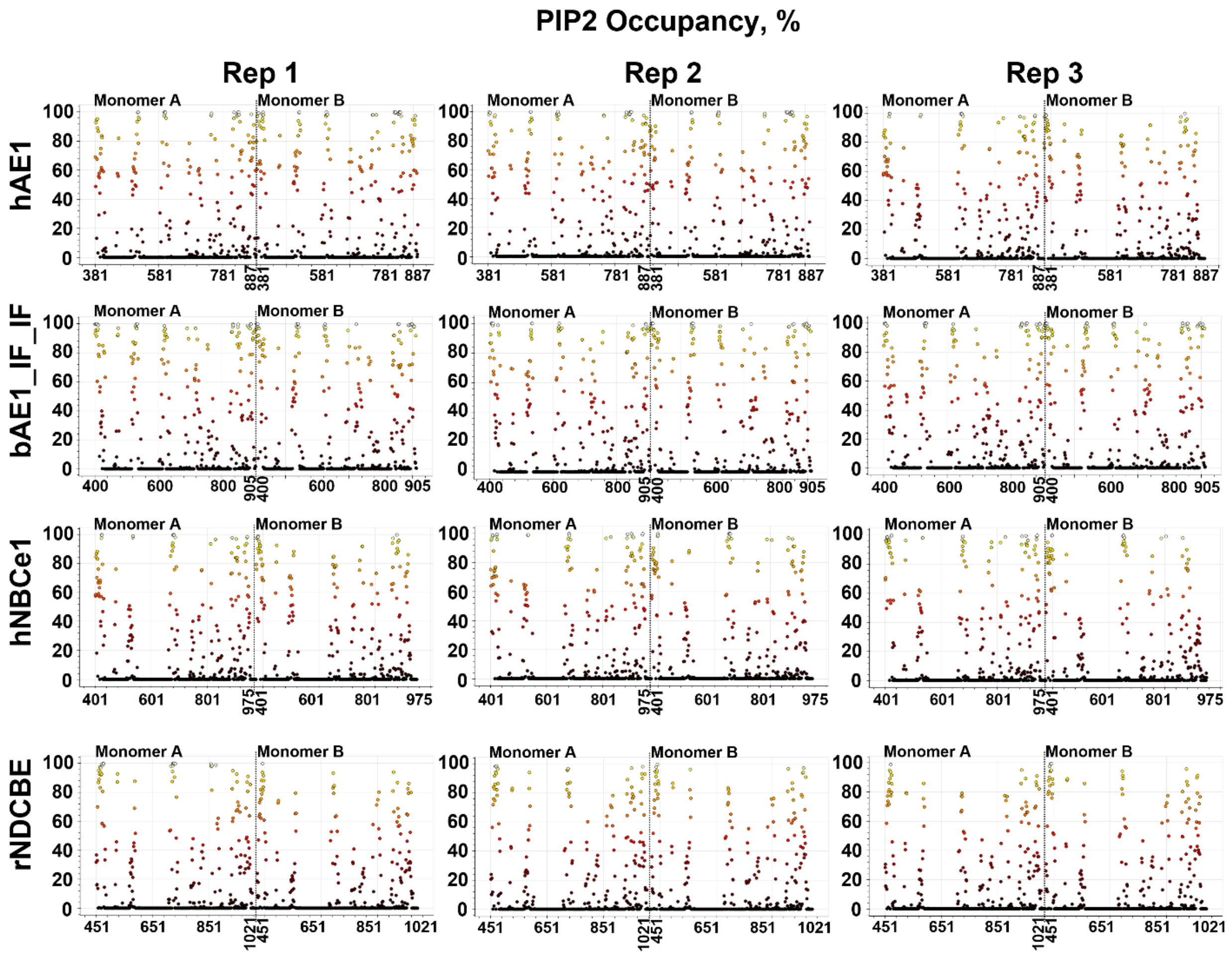


**Figure S3** Occupancy plots (in %) for PIP2-protein contacts in all simulated replicas (three replicas Rep1 – Rep3 per protein dimer). The end of the plot for the first monomer in the dimer (Monomer A) and the beginning of the plot for the second monomer of the dimer (Monomer B) is signified by a dashed line and the provided numbering of the amino acid residues along the x-axes of the plots.

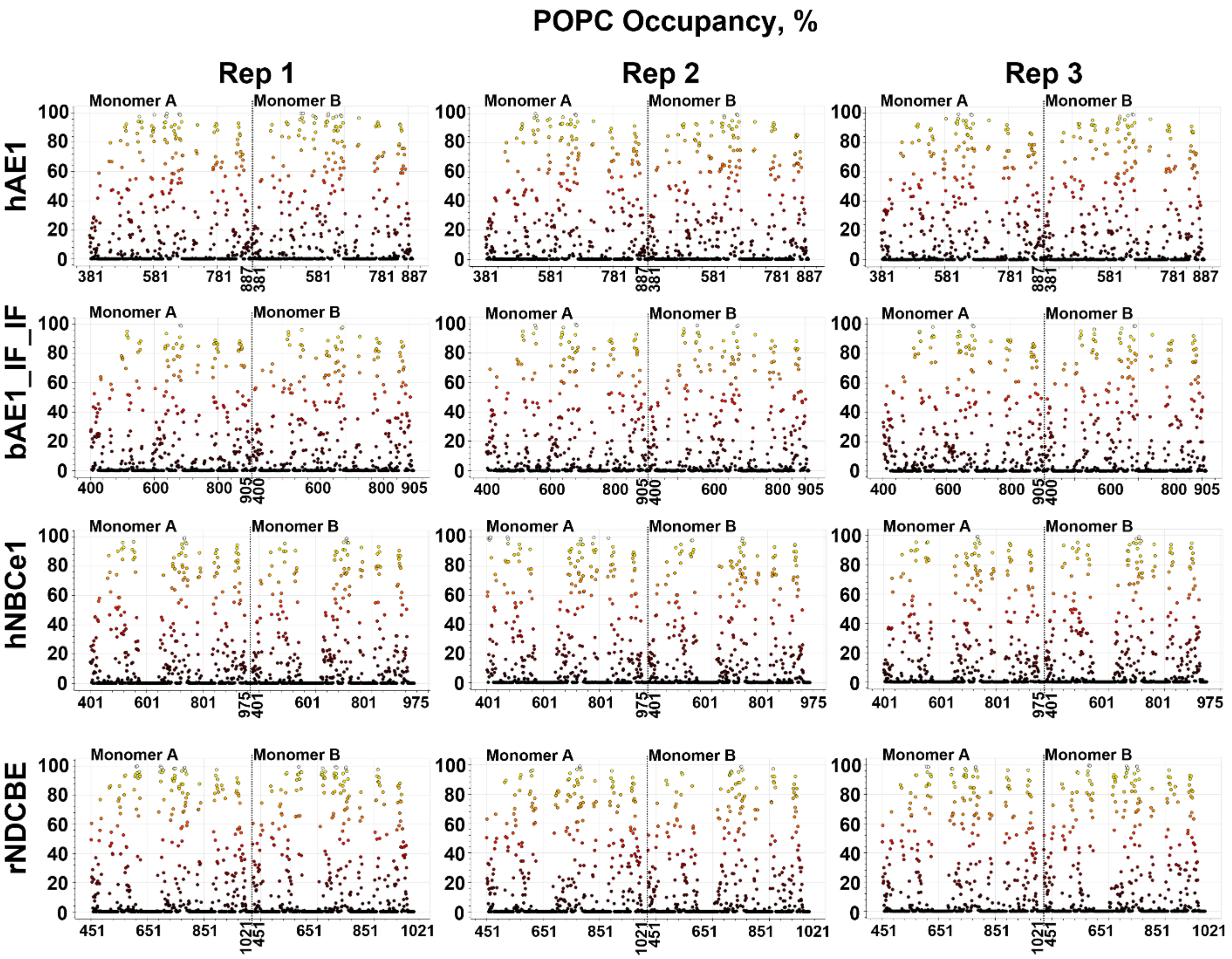


**Figure S4** Occupancy plots (in %) for POPC-protein contacts in all simulated replicas (three replicas Rep1 – Rep3 per protein dimer). The end of the plot for the first monomer in the dimer (Monomer A) and the beginning of the plot for the second monomer of the dimer (Monomer B) is signified by a dashed line and the provided numbering of the amino acid residues along the x-axes of the plots.

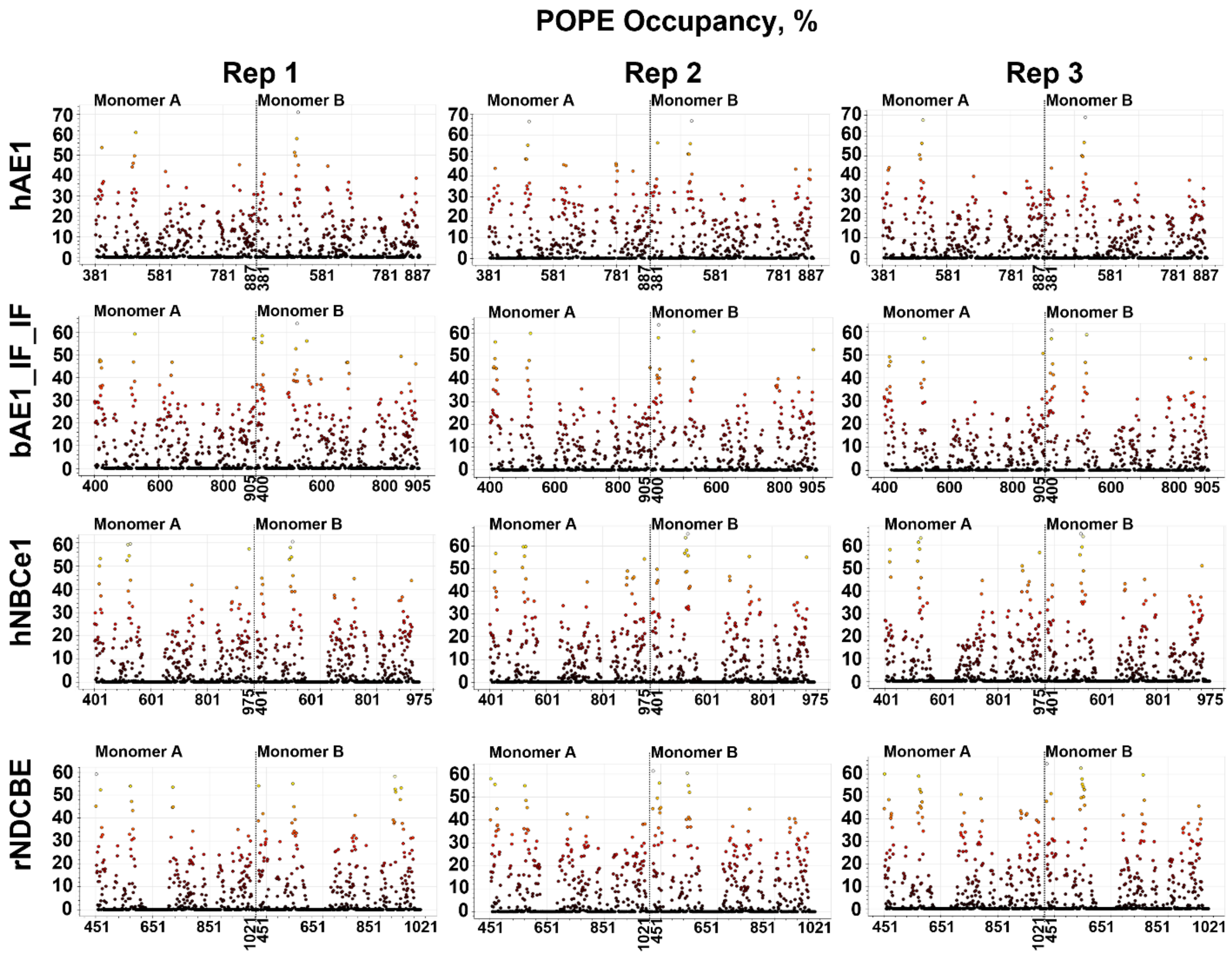


**Figure S5** Occupancy plots (in %) for POPE-protein contacts in all simulated replicas (three replicas Rep1 – Rep3 per protein dimer). The end of the plot for the first monomer in the dimer (Monomer A) and the beginning of the plot for the second monomer of the dimer (Monomer B) is signified by a dashed line and the provided numbering of the amino acid residues along the x-axes of the plots.

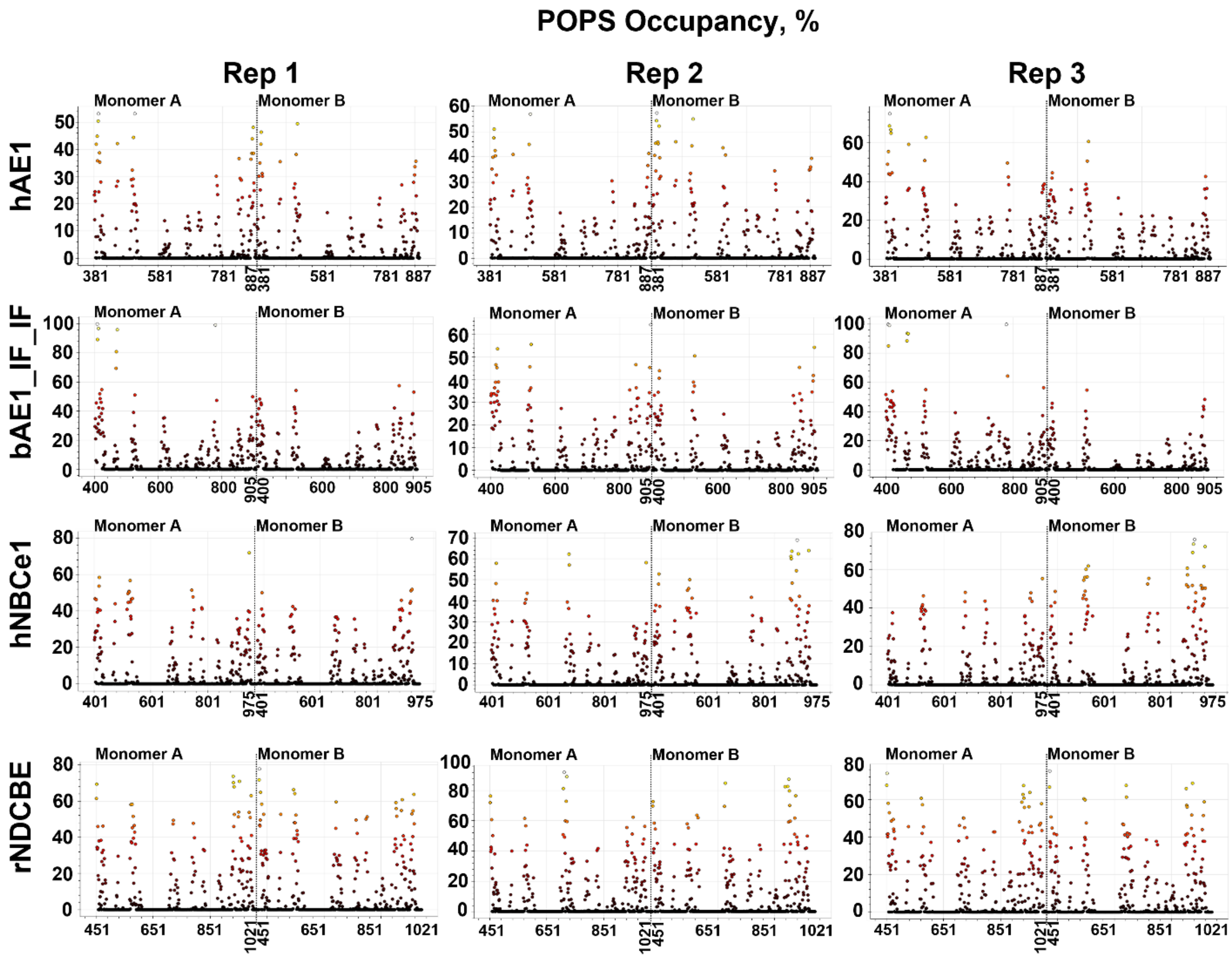


**Figure S6** Occupancy plots (in %) for POPS-protein contacts in all simulated replicas (three replicas Rep1 – Rep3 per protein dimer). The end of the plot for the first monomer in the dimer (Monomer A) and the beginning of the plot for the second monomer of the dimer (Monomer B) is signified by a dashed line and the provided numbering of the amino acid residues along the x-axes of the plots.

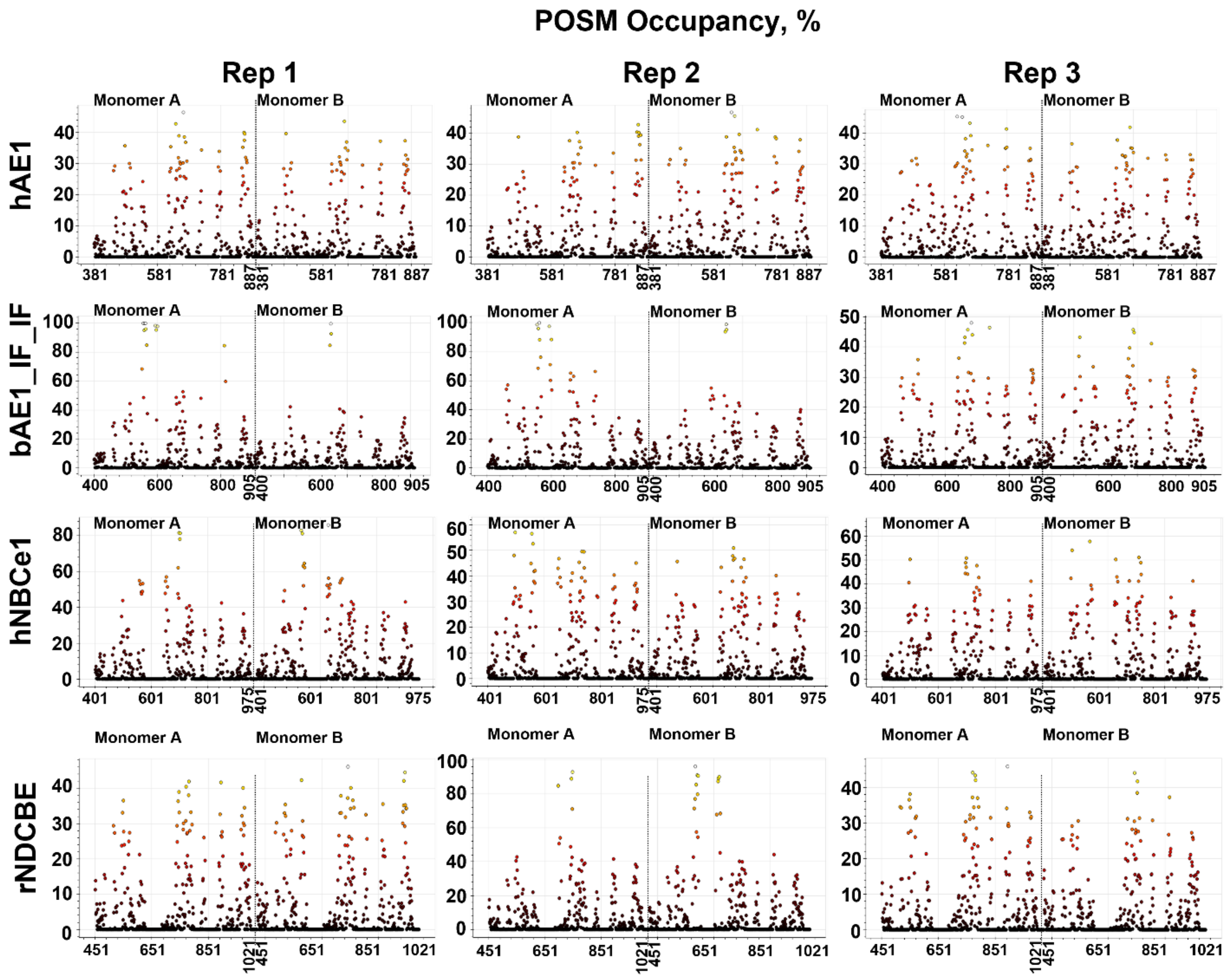


**Figure S7** Occupancy plots (in %) for POSM-protein contacts in all simulated replicas (three replicas Rep1 – Rep3 per protein dimer). The end of the plot for the first monomer in the dimer (Monomer A) and the beginning of the plot for the second monomer of the dimer (Monomer B) is signified by a dashed line and the provided numbering of the amino acid residues along the x-axes of the plots.

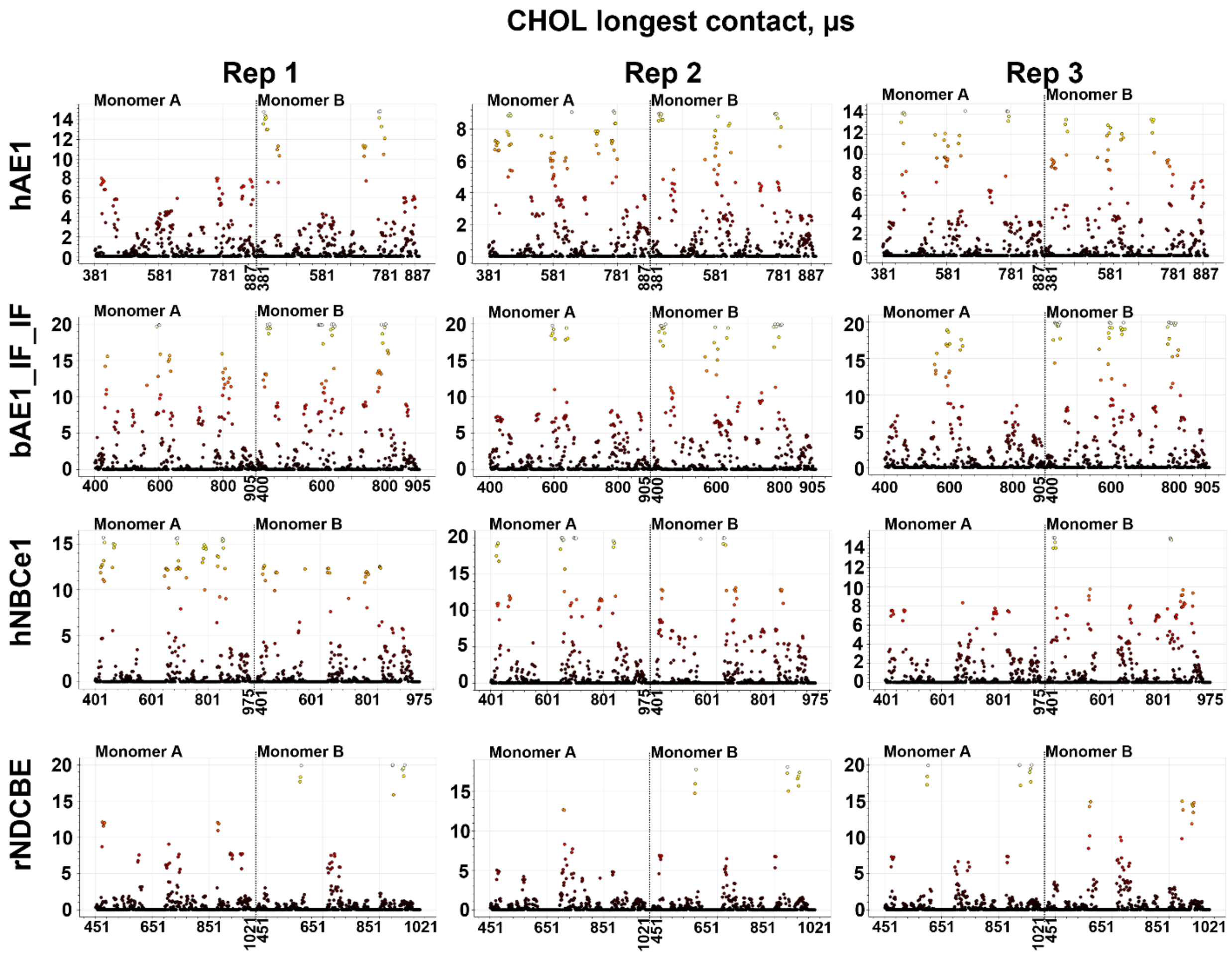


**Figure S8** Longest-contact plots (in μs) for CHOL-protein contacts in all simulated replicas (three replicas Rep1 – Rep3 per protein dimer). The end of the plot for the first monomer in the dimer (Monomer A) and the beginning of the plot for the second monomer of the dimer (Monomer B) is signified by a dashed line and the provided numbering of the amino acid residues along the x-axes of the plots.

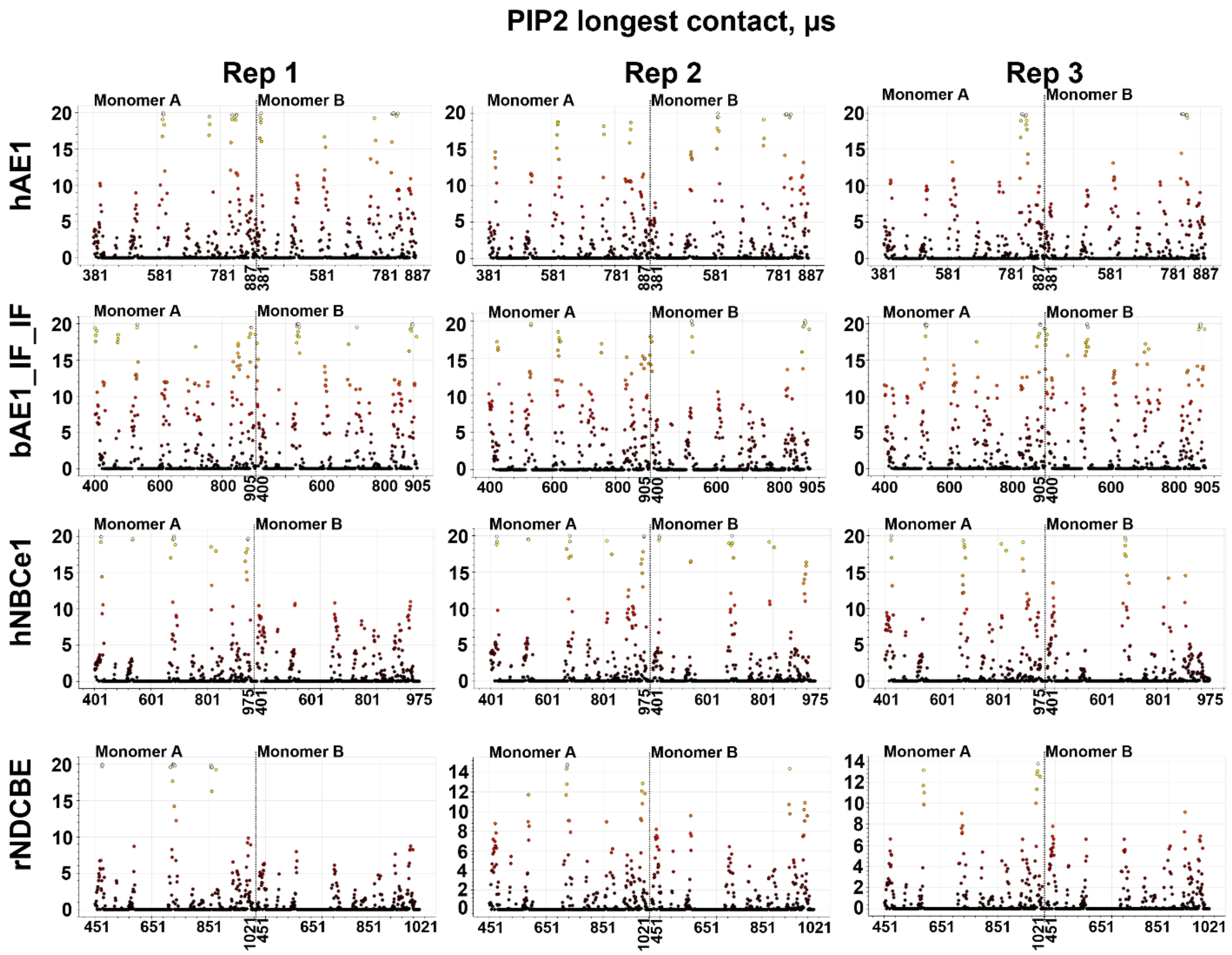


**Figure S9** Longest-contact plots (in μs) for PIP2-protein contacts in all simulated replicas (three replicas Rep1 – Rep3 per protein dimer). The end of the plot for the first monomer in the dimer (Monomer A) and the beginning of the plot for the second monomer of the dimer (Monomer B) is signified by a dashed line and the provided numbering of the amino acid residues along the x-axes of the plots.

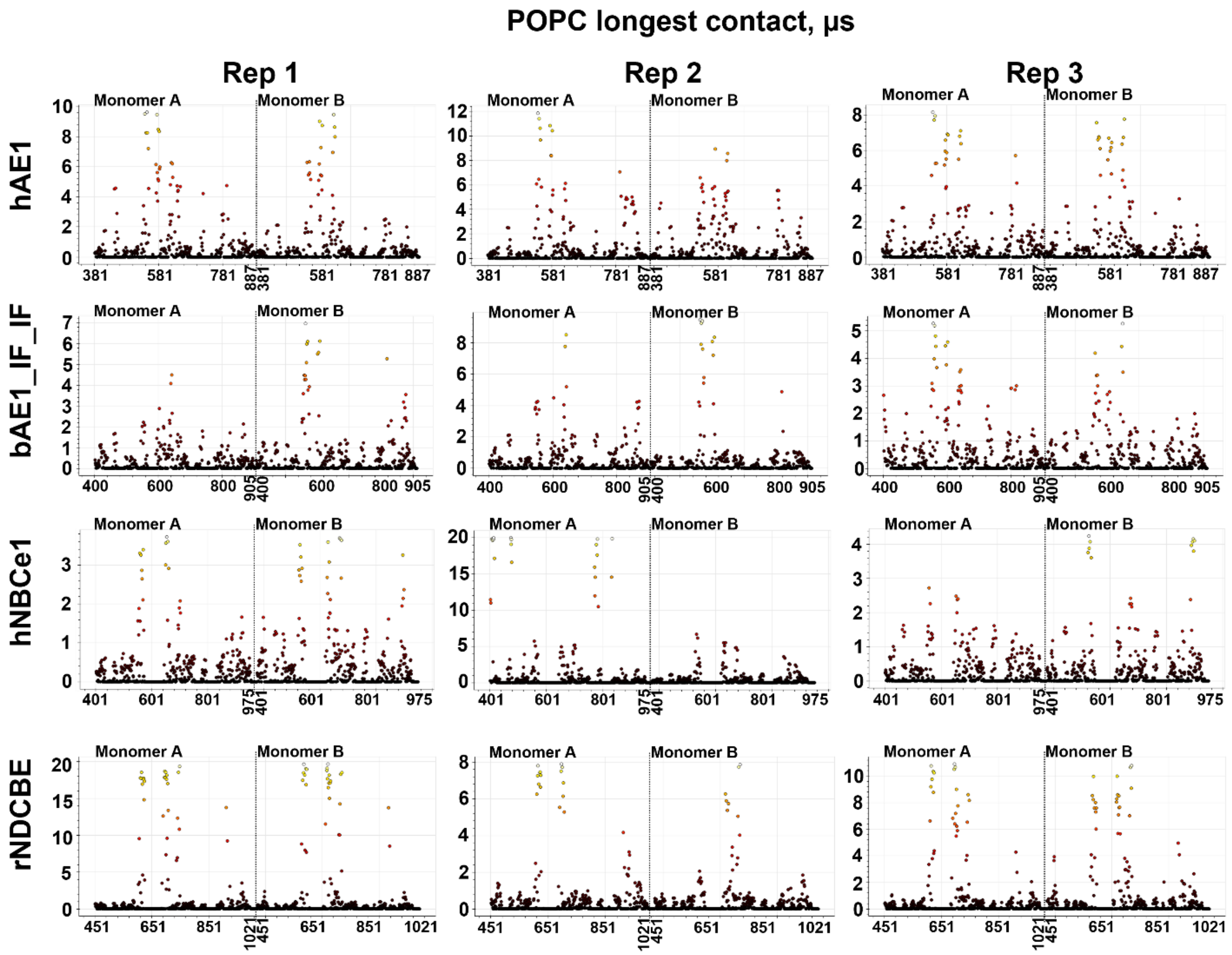


**Figure S10** Longest-contact plots (in μs) for POPC-protein contacts in all simulated replicas (three replicas Rep1 – Rep3 per protein dimer). The end of the plot for the first monomer in the dimer (Monomer A) and the beginning of the plot for the second monomer of the dimer (Monomer B) is signified by a dashed line and the provided numbering of the amino acid residues along the x-axes of the plots.

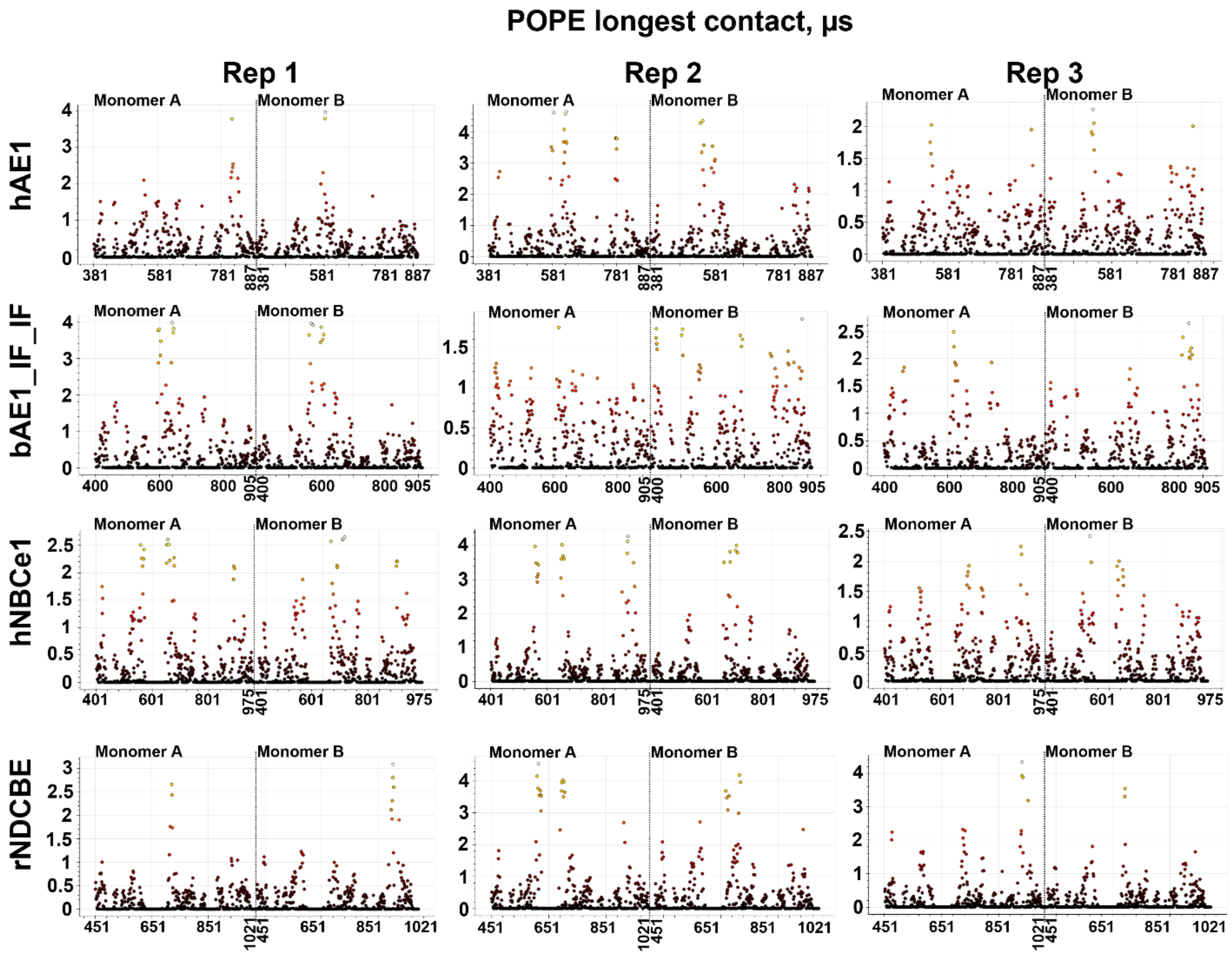


**Figure S11** Longest-contact plots (in μs) for POPE-protein contacts in all simulated replicas (three replicas Rep1 – Rep3 per protein dimer). The end of the plot for the first monomer in the dimer (Monomer A) and the beginning of the plot for the second monomer of the dimer (Monomer B) is signified by a dashed line and the provided numbering of the amino acid residues along the x-axes of the plots.

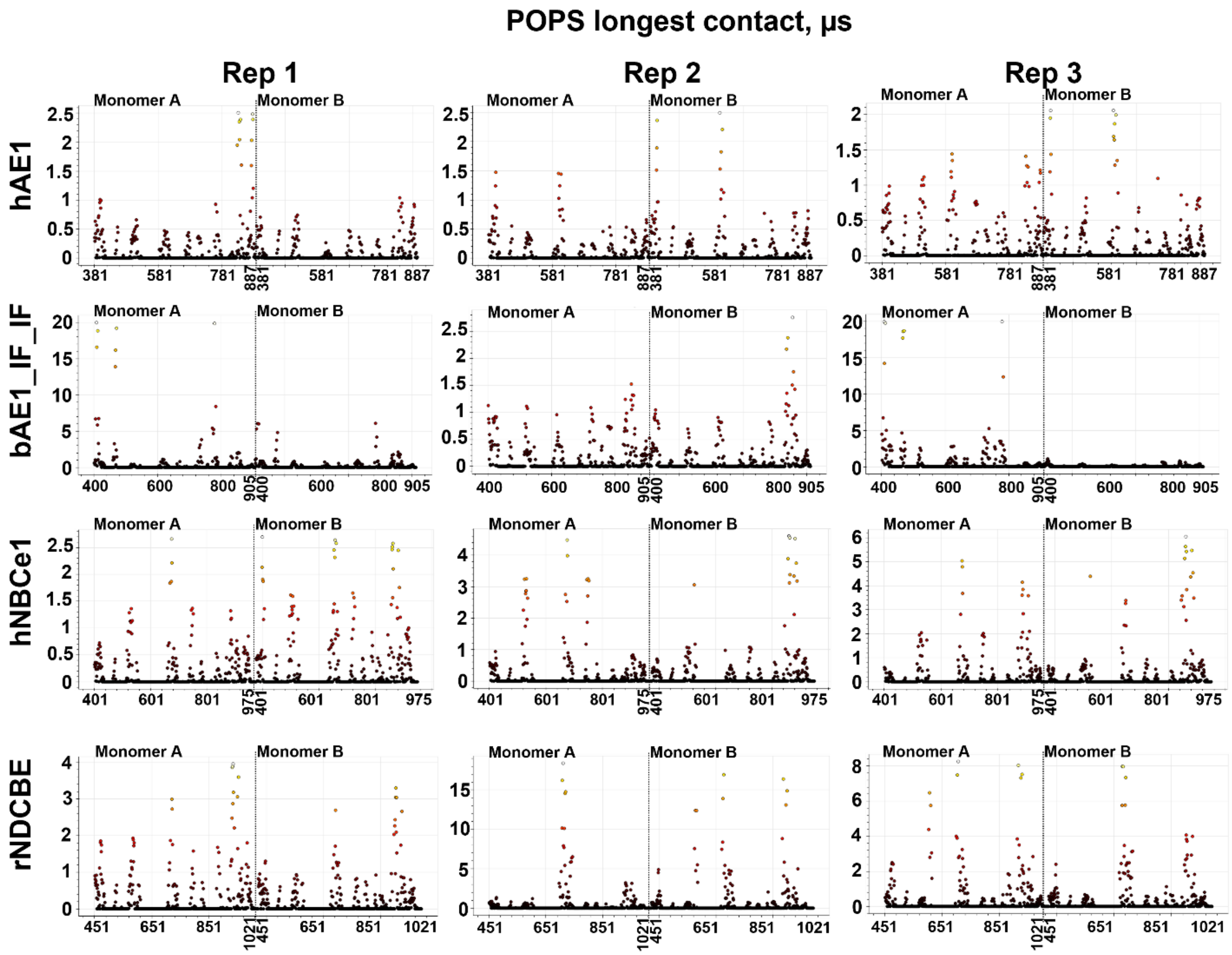


**Figure S12** Longest-contact plots (in μs) for POPS-protein contacts in all simulated replicas (three replicas Rep1 – Rep3 per protein dimer). The end of the plot for the first monomer in the dimer (Monomer A) and the beginning of the plot for the second monomer of the dimer (Monomer B) is signified by a dashed line and the provided numbering of the amino acid residues along the x-axes of the plots.

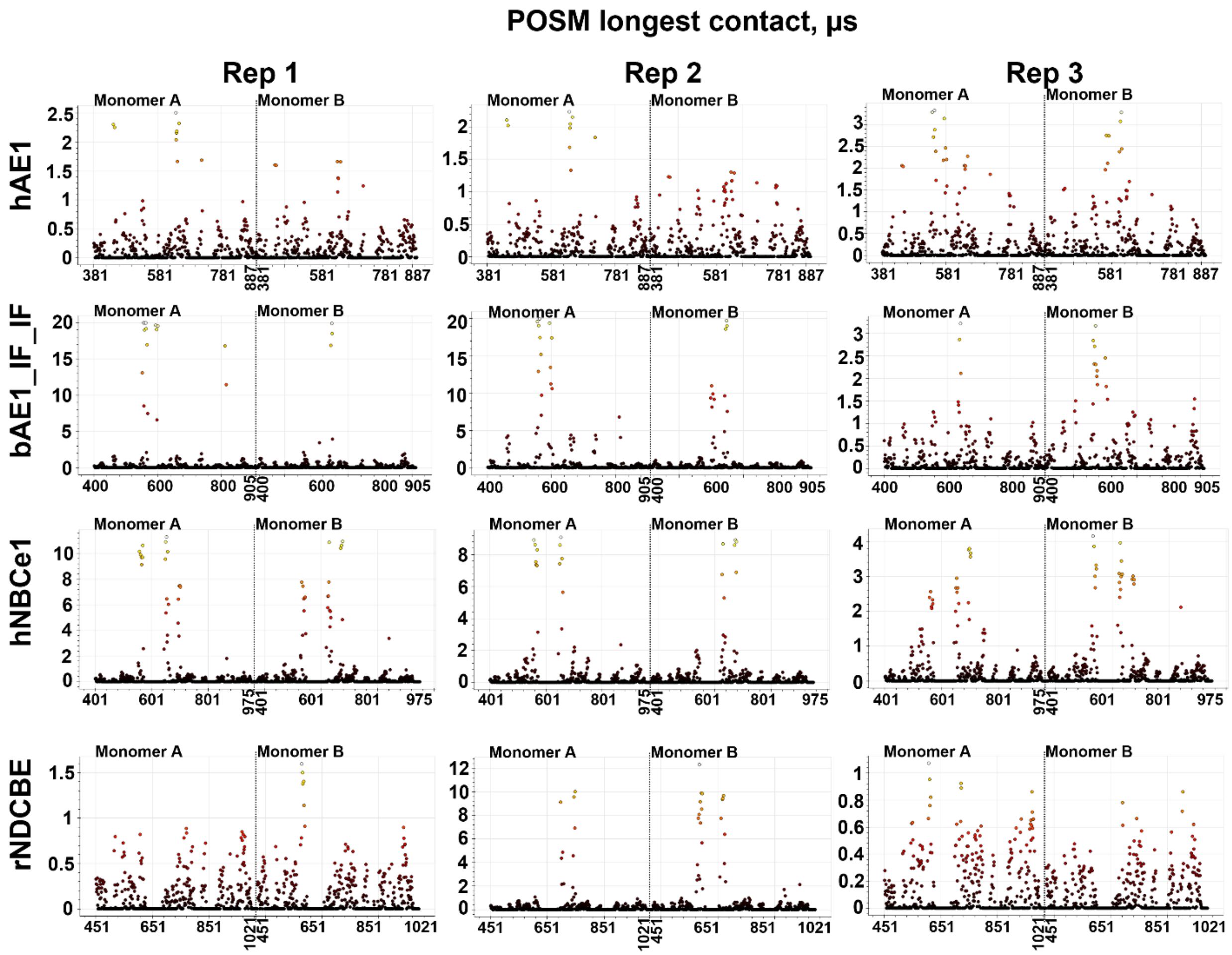


**Figure S13** Longest-contact plots (in μs) for POSM-protein contacts in all simulated replicas (three replicas Rep1 – Rep3 per protein dimer). The end of the plot for the first monomer in the dimer (Monomer A) and the beginning of the plot for the second monomer of the dimer (Monomer B) is signified by a dashed line and the provided numbering of the amino acid residues along the x-axes of the plots.

**Table S1** Comparison of the lipid compositions in the basolateral membrane of the proximal tubule (from lipidomics (1)) and model membranes of HEK293 (present work), neuron (2), erythrocyte (3), and average (2) cells.

| % lipids (no CHOL) | Proximal tubule basolateral exp | HEK293 model | Neuron model | Erythrocyte model | Average cell model |
|---|---|---|---|---|---|
| Sphingomyelins | 13.4 | 10.6 | 10.1 | 26 | 20.5 |
| Phosphatidylcholines | 38.4 | 50 | 34.0 | 31 | 38.1 |
| Phosphatidylethanolamines | 32.3 | 17.1 | 29.1 | 28 | 21.8 |
| Phosphatidylinositols | 4.31 | 4.5 | 5.9 | 2 | 4.3 |
| Phosphatidylserines | 7.5 | 17.9 | 8.8 | 13 | 7.5 |
| Others | 0.27 | 0 | 12.1 | 0 | 7.9 |
| Total % lipids (no CHOL) | 96.18 | 100.1 | 100 | 100 | 100.1 |
| CHOL/lipid ratio | 0.5 | 0.25 | 0.8 | 1 | 0.43 |

**Table S2** AE1 residues at the lipid-protein interface whose mutation impacts transport, with their corresponding locations in the three-dimensional structure of the protein and at the identified putative lipid sites from Figure 4.

| Mutated residue | Location | Lipid site | Effect |
|---|---|---|---|
| Δ400-408 (4) | H1,TM1 | $S1b_{PIP2}$, $S1_{CHOL}$ | Missfolded |
| F507 (5) | TM4 | $S2a,b_{PIP2}$ | |
| E508 (5) | H2 | $S2a,b_{PIP2}$ | |
| G509 (5) | H2 | $S2a,b_{PIP2}$ | |
| S510 (5) | H2 | $S2a,b_{PIP2}$ | |
| R518 (4) | TM5 | $S2a,b_{PIP2}$ | Spherocytosis |
| S525 (6) | TM5 | $S5_{CHOL}$ | |
| F526 (6) | TM5 | $S5_{CHOL}$ | |
| I528 (6) | TM5 | $S5_{CHOL}$ | |
| S529 (6) | TM5 | $S5_{CHOL}$ | |
| L530 (6) | TM5 | $S5_{CHOL}$ | |
| I533 (6) | TM5 | $S1_{POPC}$, $S1_{POSM}$ | |
| Y534 (6) | TM5 | $S1_{POPC}$, $S1_{POSM}$ | |
| F537 (6) | TM5 | $S1_{POPC}$, $S1_{POSM}$ | |
| L540 (6) | TM5 | $S1_{POPC}$, $S1_{POSM}$ | |
| I541 (6) | TM5 | $S1_{POPC}$, $S1_{POSM}$ | |
| P548 (4) | TM5 | $S1_{POPC}$, $S1_{POSM}$ | Blood antigen |
| F584 (5) | TM6 | $S3_{CHOL}$, $S1_{POPC}$ | |
| R602 (4) | TM7 | $S1b,c,d_{PIP2}$ | dRTA |
| I605 (5) | TM7 | $S1b,c,d_{PIP2}$ | |
| G606 (5) | TM7 | $S1b,c,d_{PIP2}$ | |
| G609 (5) | TM7 | $S4_{CHOL}$ | dRTA |
| S613 (4) | TM7 | $S4_{CHOL}$ | dRTA |
| G714 (4) | TM9 | $S2_{CHOL}$ | Spherocytosis |
| M741 (7) | LoopTM10-11 | $S1b,c_{PIP2}$ | |
| G742 (7) | LoopTM10-11 | $S1b,c_{PIP2}$ | |
| A744 (7) | LoopTM10-11 | $S1b,c_{PIP2}$ | |
| P747 (7) | LoopTM10-11 | $S1b,c_{PIP2}$ | |
| E758 (4) | TM11 | $S1,2_{CHOL}$ | Stomatocytosis |
| R760 (4,7) | TM11 | $S1,2_{CHOL}$ | Stomatocytosis |

| | | | |
|---|---|---|---|
| I761 (7) | TM11 | $S1,2_{CHOL}$ | |
| S762 (7) | TM11 | $S1,2_{CHOL}$ | |
| L765 (7) | TM11 | $S1,2_{CHOL}$ | |
| G771 (4) | TM11 | $S1,2_{CHOL}$ | Spherocytosis |
| S773 (4) | TM11 | $S1,2_{CHOL}$ | dRTA |
| I783 (4) | H3 | $S4_{CHOL}$ | Spherocytosis |
| F789 (5) | TM12 | $S4_{CHOL}$ | |
| H834 (4) | TM13 | $S1d,e_{PIP2}$ | Spherocytosis |
| T837 (4) | TM13 | $S1d,e_{PIP2}$ | Spherocytosis |
| A858 (4) | TM14 | $S2a,b_{PIP2}$ | Spherocytosis |
| P868 (4) | TM14 | $S2a,b_{PIP2}$ | $V_{max}$ increase |
| R870 (4) | TM14 | $S2a,b_{PIP2}$ | Spherocytosis |

**Table S3** NBCe1 residues at the lipid-protein interface whose mutation impacts transport, with their corresponding locations in the three-dimensional structure of the protein and at the identified putative lipid sites from Figure 4.

| Mutated residue | Location | Lipid site | Effect |
|---|---|---|---|
| D405 (8) | H1 | $S3a,b_{PIP2}$, $S1a_{PIP2}$ | |
| L421 (8) | H1,TM1 | $S1b_{PIP2}$, $S1_{CHOL}$ | |
| N422 (8) | H1,TM1 | $S1b_{PIP2}$, $S1_{CHOL}$ | |
| A425 (9) | TM1 | $S1b_{PIP2}$, $S1_{CHOL}$ | |
| S427 (8,9) | TM1 | $S1b_{PIP2}$, $S1_{CHOL}$ | pRTA |
| I429 (9) | TM1 | $S1b_{PIP2}$, $S1_{CHOL}$ | |
| Y433 (9) | TM1 | $S1_{CHOL}$ | |
| T436 (9) | TM1 | $S1_{CHOL}$, $S4_{CHOL}$ | |
| T438 (9) | TM1 | $S1_{CHOL}$, $S4_{CHOL}$ | |
| R538 (10) | H2 | $S2a,b_{PIP2}$ | |
| K558 (11) | TM5 | $S1_{POPC}$, $S1_{POSM}$ | |
| D647 (10) | TM6 | $S3_{CHOL}$, $S1_{POPC}$, $S1_{POSM}$ | |
| F656 (10) | TM6 | $S3_{CHOL}$, $S1_{POPC}$, $S1_{POSM}$ | |
| T677 (10) | TM7 | $S1b,c,d_{PIP2}$ | |
| R680 (10) | TM7 | $S1b,c,d_{PIP2}$ | |
| K681 (10) | TM7 | $S1b,c,d_{PIP2}$ | |
| K854 (10) | H3 | $S4_{CHOL}$ | |

| | | | |
|---|---|---|---|
| R904 (10) | TM13 | S1d,$e_{PIP2}$ | |
| R905 (10) | TM13 | S1d,$e_{PIP2}$ | |
| H907 (10) | TM13 | S1d,$e_{PIP2}$ | |
| Q913 (12) | TM13 | S1d,$e_{PIP2}$, $S5_{CHOL}$ | pRTA |
| R943 (10) | TM14 | S2a,$b_{PIP2}$ | |

**Table S4** CRAC/CARC-like domains identified in the sequences of the four studies systems from ScanProsite (13) sequence analysis with their corresponding locations in the three-dimensional structure of the protein and at the identified putative lipid sites from Figure 4.

| CRAC domains | | | |
|---|---|---|---|
| Protein | Domain sequence | Location | Lipid sites |
| hAE1 | L484-R490 | TM4 | N/A |
| | L530-K539 | TM5 | $S5_{CHOL}$, $S1_{POPC}$, $S1_{POSM}$ |
| | V822-K829 | H5 | S1d,$e_{PIP2}$ |
| bAE1 | L502-R508 | TM4 | N/A |
| | L548-K557 | TM5 | $S5_{CHOL}$, $S1_{POPC}$, $S1_{POSM}$ |
| | L641-K649 | TM7 | $S3_{CHOL}$, $S1_{POPC}$, $S1_{POSM}$ |
| | V840-K847 | H5 | S1d,$e_{PIP2}$ |
| hNBCe1 | L532-R538 | H2 | S2a,$b_{PIP2}$ |
| rNDCBE | L557-R563 | TM4 | N/A |
| | L585-R591 | H2 | S2a,$b_{PIP2}$ |
| | L628-R637 | LoopTM5-6 | N/A |
| | L649-K653 | LoopTM5-6 | N/A |
| CARC domains | | | |
| hAE1 | R384-L394 | H1 | S3a,$b_{PIP2}$, $S1a_{PIP2}$ |
| | R490-V501 | TM4 | N/A |
| | R518-L530 | TM5 | $S5_{CHOL}$ |
| | K539-L549 | TM5 | $S1_{POPC}$, $S1_{POSM}$ |
| | K551-V560 | LoopTM5-6 | N/A |
| | K590-L601 | TM6-TM7 | S1b,c,$d_{PIP2}$ |
| | R602-V610 | TM7 | $S4_{CHOL}$, S1b,c,$d_{PIP2}$ |
| | K698-L708 | LoopTM8-9 | N/A |
| | K814-V822 | LoopH4-H5 | $S1d_{PIP2}$ |
| | R901-V907 | Ct | N/A |

| | | | |
|---|---|---|---|
| | R402-L412 | H1 | S3a,b$_{PIP2}$, S1a$_{PIP2}$ |
| | R508-V519 | TM4 | N/A |
| | R536-L548 | TM5 | S5$_{CHOL}$ |
| | K557-L567 | TM5 | S1$_{POPC}$, S1$_{POSM}$ |
| | K569-L576 | LoopTM5-6 | N/A |
| bAE1 | K608-L619 | TM6-TM7 | S1b,c,d$_{PIP2}$ |
| | R620-V628 | TM7 | S4$_{CHOL}$, S1b,c,d$_{PIP2}$ |
| | K716-L726 | LoopTM8-9 | N/A |
| | K832-V840 | LoopH4-H5 | S1d$_{PIP2}$ |
| | R850-V860 | TM13 | S5$_{CHOL}$, S1d,e$_{PIP2}$ |
| | R889-L893 | LoopTM14-H6 | S2a,b$_{PIP2}$ |
| | K500-L511 | TM3-TM4 | N/A |
| hNBCe1 | R538-L547 | TM5 | S5$_{CHOL}$ |
| | K574-L580 | LoopTM5-6 | N/A |
| | K627-V638 | LoopTM5-6 | N/A |
| | R680-L690 | TM7 | S4$_{CHOL}$, S1b,c,d$_{PIP2}$ |
| | K770-L779 | LoopTM8-9 | N/A |
| | K890-V901 | LoopH4-H5 | S1d$_{PIP2}$ |
| hNBCe1 | R904-L915 | TM13 | S5$_{CHOL}$, S1d,e$_{PIP2}$ |
| | R905-L917 | TM13 | S5$_{CHOL}$, S1d,e$_{PIP2}$ |
| | K944-L955 | LoopTM14-H6 | S2a,b$_{PIP2}$ |
| | K546-L557 | TM3 | N/A |
| | K550-L560 | TM3 | N/A |
| | K553-L562 | TM3-TM4 | N/A |
| | R591-L600 | TM5 | S5$_{CHOL}$ |
| rNDCBE | K716-V725 | TM6-TM7 | S1b,c,d$_{PIP2}$ |
| | R726-L736 | TM7 | S4$_{CHOL}$, S1b,c,d$_{PIP2}$ |
| | K816-V827 | LoopTM8-9 | N/A |
| | K936-V947 | LoopH4-H5 | S1d$_{PIP2}$ |
| | R950-L960 | TM13 | S5$_{CHOL}$, S1d,e$_{PIP2}$ |
| | K990-L1001 | LoopTM14-H6 | S2a,b$_{PIP2}$ |
